# Sequence architecture of RNA reveals key molecular signatures driving RNA phase separation

**DOI:** 10.64898/2026.10.23.761066

**Authors:** Sonali Chatterjee, Ranjit Prasad Bahadur

## Abstract

Biomolecular condensates are dynamic membraneless assemblies that organize various cellular processes through liquid–liquid phase separation (LLPS). RNA is an active regulator of biomolecular condensates and influence their formation through sequence-encoded features. Despite its established role in condensate formation, no computational framework exists to identify underlying signatures to predict LLPS-associated RNAs. We develop an interpretable hybrid framework to distinguish reviewed, experimentally supported LLPS-associated RNAs from background RNAs. We uncover biologically meaningful sequence properties associated with condensate participation. Each RNA is represented by 167 handcrafted descriptors capturing composition, k-mer usage, sequence complexity, repeat organization and dinucleotide autocorrelation together with 640-dimensional contextual embeddings from RNA-FM. These representations are integrated through cross-validated stacking with neural-network meta-learner. The framework achieves an AUC-ROC of 0.935, AUC-PR of 0.944 and MCC of 0.782. Feature statistics, ablation analysis and SHAP interpretation reveal a distributed sequence architecture dominated by altered local k-mer usage with additional contributions from low-complexity burden, repeat-associated organization and long-range dinucleotide pattern. Biological case studies including MALAT1 and TMPO-AS1 further connect these patterns with experimentally characterized modes of multivalent RNA–protein assembly. Together, the results show how interpretable machine learning can move beyond prediction to identify sequence features that generate mechanistic hypotheses for RNA-mediated condensation.

## 1. Introduction

Biomolecular condensates formed through liquid–liquid phase separation (LLPS) organize cellular processes by concentrating proteins, nucleic acids and regulatory factors without membrane-bound compartments [1–4]. Rather than a passive passenger, RNA is increasingly recognized as an active contributor to condensate formation [5–8]. Condensates such as nucleoli, stress granules, P-bodies, paraspeckles and DNA-damage-associated assemblies are enriched for particular RNA species, including lncRNAs, mRNAs, rRNAs, eRNAs and satellite RNAs [5–9]. Depending on context, RNAs may act as scaffolds, clients, regulators of protein phase separation or spatial organizers of ribonucleoprotein assemblies [5–8]. These roles are likely shaped by multiple layers of RNA sequence organization, including local composition, short k-mer grammar, low-complexity or repetitive regions, motif spacing and long-range sequence-order effects [5–10]. Together, these properties may support RNA–protein recognition, RNA–RNA contacts, local structural preferences and multivalent interactions central to condensate biology [5, 7–8, 10]. Aberrant RNA expression, localization, modification or folding can alter condensate assembly, composition and physical properties [9]. Dysregulation of RNA-mediated condensate behavior has been linked to cancer and neurodegeneration [11]. For example, repeat-expansion RNAs, such as expanded CAG or G4C2 repeat transcripts, can sequester RNA-binding proteins and perturb RNA processing pathways by forming aberrant RNA-protein condensates [10–11]. Altered RNA–protein condensate interactions are also associated with pathological aggregation of RNA-binding proteins (RBP) such as TDP-43 and FUS [2, 4, 9, 11]. These observations highlight the need to identify RNA sequence features associated with condensate participation and dysregulation.

Despite growing experimental evidence for the roles of RNA in condensate biology, computational framework for sequence-based prediction of LLPS-associated RNAs yet to be developed. Existing computational tools have focused exclusively on proteins, using amino acid composition, intrinsically disordered regions and prion-like domain features that have no direct RNA equivalent [12–17]. Whether experimentally supported LLPS-associated RNAs exhibit reproducible primary-sequence signatures that distinguish them from LLPS-unannotated RNAs remains an open question. Curated resources such as RPS 2.0 now provide experimentally supported LLPS-associated RNA annotations [18], creating an opportunity to examine this question systematically. A primary-sequence-based approach is particularly attractive because RNA sequences are widely available, whereas experimentally resolved structures, interaction maps and condition-specific conformations remain unavailable for most transcripts.

In this study, we investigate molecular signatures for LLPS-associated RNAs that distinguish them from LLPS-unannotated background RNAs. We further ask whether these signatures reveal broader principles of RNA sequence organization relevant to condensate participation. To address this question, we use interpretable machine learning not only as a predictive framework, but also as a systematic approach for identifying and prioritizing biologically meaningful sequence features. We integrate 167 handcrafted descriptors capturing k-mer usage, low-complexity burden, repeat organization, dinucleotide autocorrelation and predicted secondary-structure properties with 640-dimensional contextual embeddings from RNA-FM [19–20]. These complementary representations are combined in a cross-validated stacking framework with a neural-network meta-learner. Beyond classification, feature-level statistics, representation ablation and SHAP interpretation are used to determine sequence signatures distinguishing LLPS-associated RNAs. The analyses reveal sequence architecture dominated by altered local k-mer usage with additional contributions from low-complexity burden, repeat-associated organization, long-range dinucleotide pattern and learned RNA sequence context. Together, this framework uses machine learning as a tool to connect predictive sequence patterns with biologically interpretable hypotheses about multivalent RNA–protein interactions and condensate-associated RNA behaviour.

## 2. Materials and Methods

### 2.1 Dataset construction and benchmark definition

Experimentally supported LLPS-associated RNAs were obtained from the reviewed subset of RPS 2.0 and designated LLPS+. Background RNA sequences were collected from Ensembl and RNAcentral [18, 21–22]. Sequences matching reviewed or high-throughput-supported RPS 2.0 entries were removed from the background pool, and the remaining sequences were designated LLPS−. Sequence redundancy was reduced on the combined LLPS+ and LLPS− dataset using CD-HIT-EST at an 85% sequence-identity threshold [23–24]. The final benchmark comprised 915 RNAs, divided into a training set of 732 sequences and a test set of 183 sequences (Table 1). Model development, preprocessing and cross-validation were restricted to the training set. The overall dataset construction and benchmark-definition workflow is summarized in Figure 1A.

**Figure 1.**
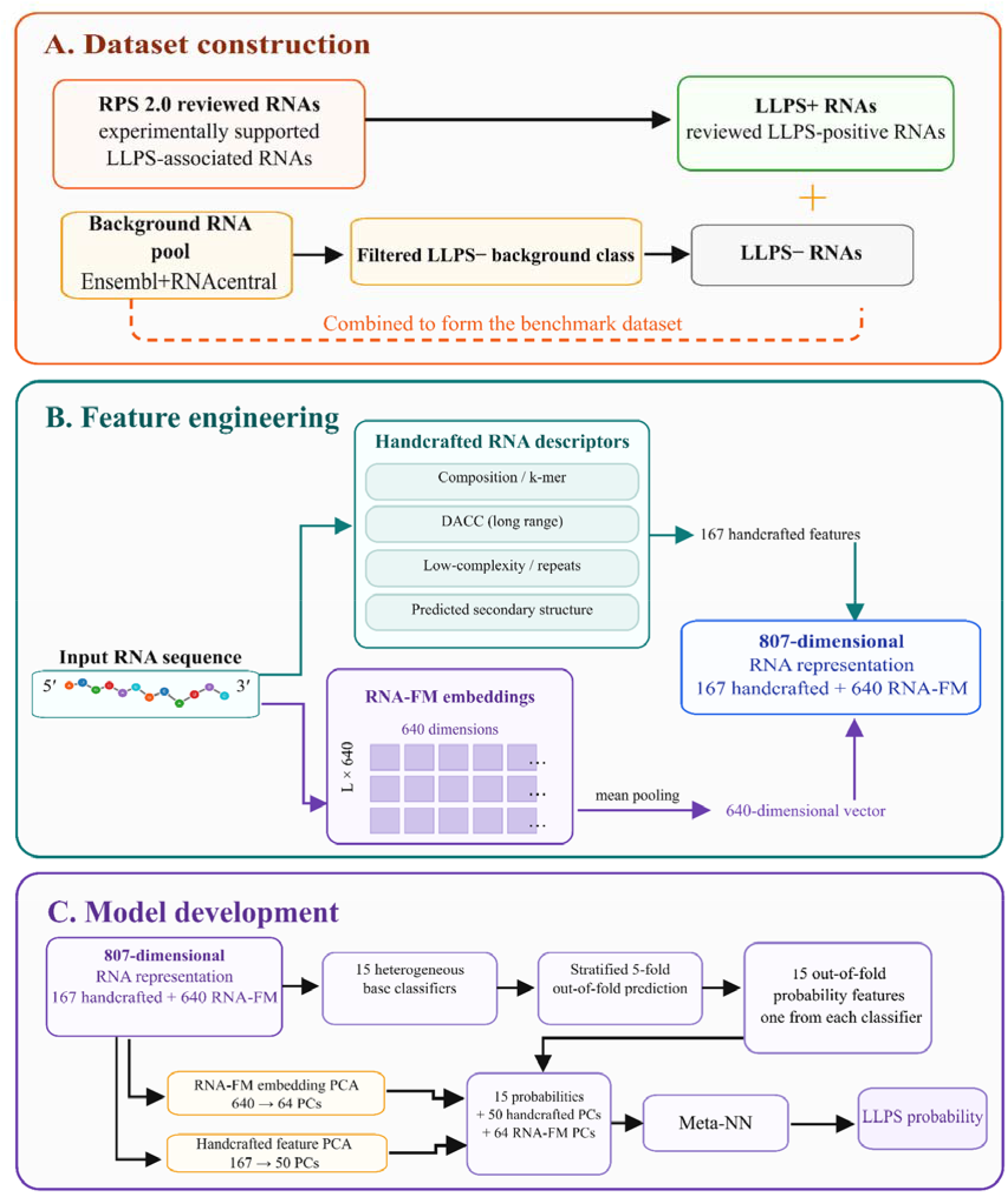
Dataset construction, feature engineering and stacked model workflow. (A) Reviewed LLPS-associated RNAs from RPS 2.0 are used as the LLPS+ class. Background RNAs from Ensembl and RNAcentral are filtered to remove known RPS 2.0 reviewed and high-throughput-supported LLPS-associated RNAs, generating the LLPS− background class. (B) RNA sequence is represented using 167 handcrafted descriptors and 640 mean-pooled RNA-FM embedding dimensions, producing an 807-dimensional RNA representation. (C) The stacked model combined stratified five-fold out-of-fold probabilities from 15 base classifiers with separately PCA-compressed handcrafted and RNA-FM feature blocks. These inputs are integrated by a Meta-NN to generate the final LLPS+ probability.

**Table 1.**
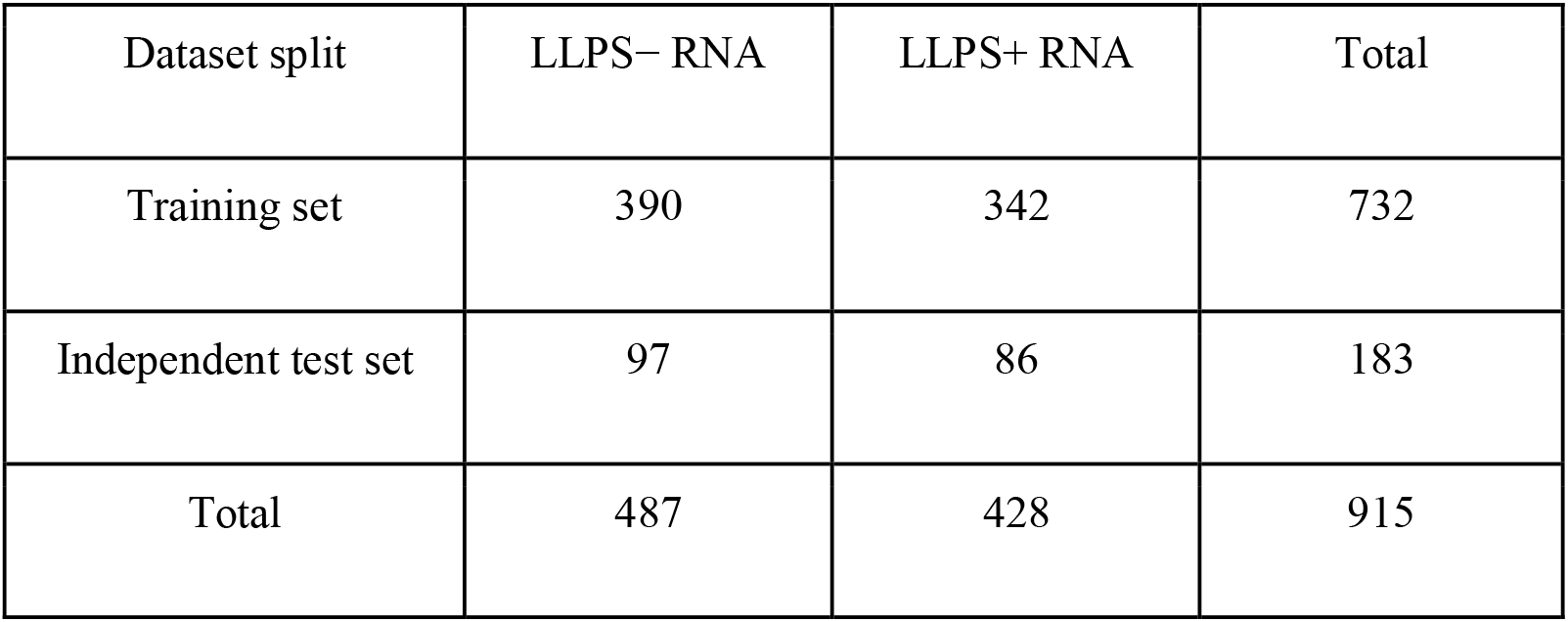
Composition of the LLPS-associated RNA benchmark dataset.

| Dataset split | LLPS− RNA | LLPS+ RNA | Total |
| --- | --- | --- | --- |
| Training set | 390 | 342 | 732 |
| Independent test set | 97 | 86 | 183 |
| Total | 487 | 428 | 915 |

### 2.2 Handcrafted RNA sequence descriptors

#### 2.2.1 Composition, k-mer and DACC descriptors

Nucleotide-composition, di- and trinucleotide-frequency and dinucleotide auto-cross-covariance (DACC) descriptors were calculated using iLearnPlus [25–26]. Composition and k-mer features represented local nucleotide (nt) usage and short sequence-word patterns, whereas DACC descriptors captured correlations between dinucleotide physicochemical properties across sequence positions, providing information on long-range sequence order and spacing.

#### 2.2.2 Custom RNA sequence complexity and repeat descriptors

Before feature calculation, sequences were converted to uppercase, thymine was replaced with uracil and noncanonical nucleotides were removed. Custom descriptors were calculated to represent sequence entropy, linguistic complexity, nucleotide-level low-complexity burden, compressibility, homopolymer organization, tandem repeats and motif clustering [27–30]. Sequence entropy was calculated from mono-, di- and trinucleotide frequency distributions as

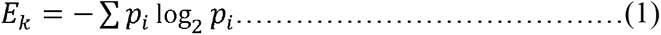

here E_k_ represents the entropy for k-mers of length k and pi is the observed frequency of the ith k-mer. Linguistic complexity was calculated as the mean observed-to-possible k-mer ratio across k-mer lengths from 1 to 5 as

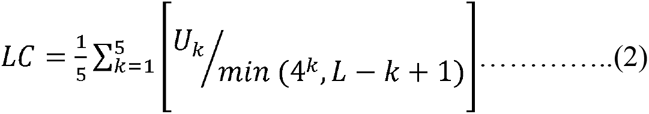

where U_k_ is the number of unique k-mers observed at length k, L is the RNA sequence length, 4□ is the total number of theoretically possible RNA k-mers of length k, and (L − k + 1) is the number of overlapping k-mer positions available in the sequence. Nucleotide-level low-complexity burden was calculated using overlapping 12-nt windows based on entropy and nucleotide diversity. Additional descriptors represented gzip compressibility, nucleotide-specific homopolymer runs, sequence coverage by runs of length ≥4 and tandem repeats of motif lengths 1–3 nt requiring at least three consecutive repeat units. Motif density and clustering were calculated using 30-nt sliding windows. Complete definitions and equations for all custom descriptors are provided in Supplementary Table S1.

#### 2.2.3 Predicted secondary-structure descriptors

Minimum-free-energy of RNA secondary structures were predicted using ViennaRNA through the RNA fold-compound interface [31]. Extracted descriptors included minimum free energy, length-normalized minimum free energy, paired and unpaired fractions, stem count, maximum stem length and mean stem length. Paired and unpaired fractions were calculated from the predicted dot-bracket structure, and stem statistics were derived from contiguous paired regions. The feature families and corresponding computational tools are summarized in Supplementary Table S2.

### 2.3 Generation of RNA-FM embedding

To complement the handcrafted descriptors, each RNA sequence was represented using RNA-FM embeddings [19–20]. RNA-FM is a pretrained RNA language model that learns latent representations from large-scale RNA sequence data using self-supervised learning. The model is trained on approximately 23.7 million non-coding RNA sequences and has been shown to encode information relevant to RNA structure and function [19–20]. RNA-FM generates a contextual embedding matrix for each input sequence, where a sequence of length L is represented as an L × 640 matrix. Thus, each nucleotide position is represented by a 640-dimensional contextual embedding. To obtain one fixed-length representation per RNA, residue-level embeddings were mean-pooled across the sequence dimension. Sequences exceeding the model input length were divided into overlapping 512-nt windows with a stride of 256 nt. Embeddings were generated independently for each window, mean-pooled across nucleotide positions and averaged across windows to obtain one 640-dimensional representation for the complete RNA. The final mean-pooled RNA-FM vector was used as a deep sequence representation complementary to handcrafted features. RNA-FM embeddings were treated as a separate feature block during downstream modelling.

### 2.4 Feature matrix construction

For individual RNA sequence, handcrafted descriptors and RNA-FM embeddings were assembled into numerical feature matrices. Redundancy among handcrafted descriptors was reduced using pairwise Spearman correlation analysis. For feature pairs with ∣ρ∣≥0.90, one descriptor was removed. This filtering was applied only to the handcrafted feature block, retaining 167 descriptors comprising composition and k-mer features, DACC descriptors, sequence-complexity, repeat features and predicted secondary-structure properties. These were combined with 640 mean-pooled RNA-FM dimensions to produce an 807-dimensional representation for each RNA (Table 2). Class labels were encoded as LLPS+ for reviewed RPS 2.0 RNAs and LLPS− for filtered background RNAs [18]. The handcrafted and RNA-FM features were retained as separate blocks for block-wise dimensionality reduction and ablation analysis. The feature-engineering workflow is shown in Figure 1B.

**Table 2.**
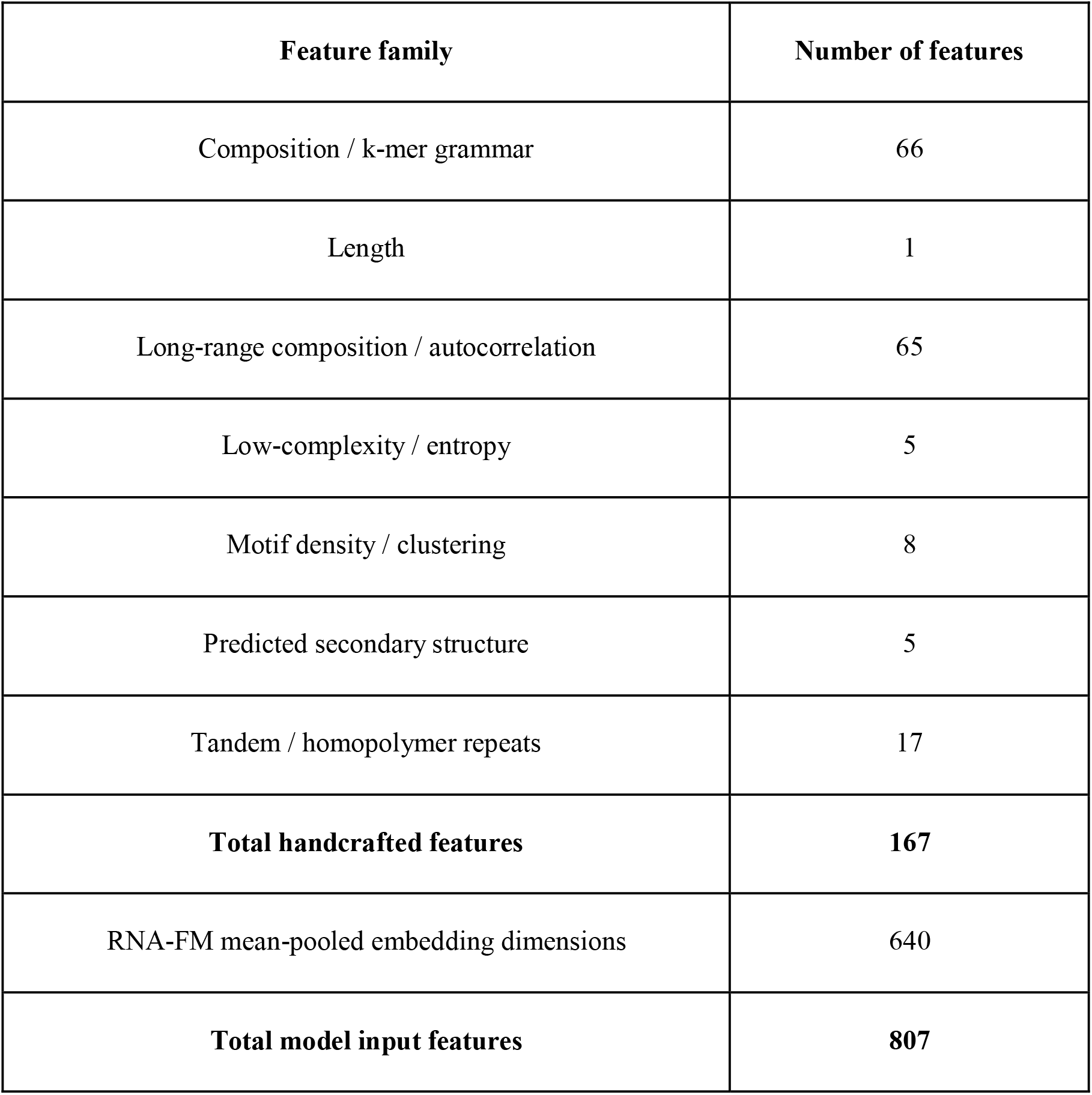
RNA sequence-derived feature representation. Individual RNA is encoded using 167 handcrafted descriptors and 640 RNA-FM embedding dimensions.

### 2.5 Statistical analysis of handcrafted RNA features

Handcrafted descriptors were compared between LLPS+ and LLPS− RNAs using two-sided Mann–Whitney U tests [32]. P-values were adjusted for multiple testing using the Benjamini–Hochberg false-discovery-rate procedure [33]. Effect sizes were calculated as rank-biserial correlations with positive values indicating higher values in LLPS+ RNAs and negative values indicating higher values in LLPS− RNAs [34]. All statistical comparisons were performed on the training set only (LLPS− = 390; LLPS+ = 342); held-out test RNAs were excluded to prevent information leakage. For each descriptor, class-wise mean and median, raw p-values, FDR-adjusted q-values, effect sizes and directions of change were reported. Features satisfying both the significance criterion of q < 0.05 and the predefined effect-size threshold of |r_rb| ≥ 0.147 were considered robustly class-separating. Complete results are provided in Supplementary Table S3.

### 2.6 Stacked machine-learning framework

#### 2.6.1 Base classifiers and out-of-fold prediction

To capture complementary decision boundaries, we trained a heterogeneous base layer of 15 classifiers: logistic regression with L2 regularization [35–36], ridge classifier [36], linear discriminant analysis [37], k-nearest neighbours [38], Gaussian naïve Bayes [39], random forest [40], extra trees [41], histogram gradient boosting and gradient boosting [42], AdaBoost [43], LightGBM [44], XGBoost [45], CatBoost [46], RBF-SVM [47], and a multilayer perceptron (MLP) [48]. Conventional classifiers were implemented in scikit-learn; XGBoost, LightGBM and CatBoost via their respective libraries [44–46, 49]. Base-model class probabilities were obtained by stratified five-fold out-of-fold prediction [50]. In each fold, the model was trained on four-fifths of the data and predicted on the held-out fifth, yielding one probability column per classifier across all training sequences. Base models were refitted on the full training set to generate test-set predictions. The resulting 15-column probability matrices served as stacking features for the meta-learner (Figure 1C).

#### 2.6.2 Meta-learner construction and training

The meta-learner integrates base-model outputs with compressed representations of both feature blocks. Using principal component analysis (PCA) [51], 167-dimensional handcrafted block was standardized and reduced to 50 PCs, and 640-dimensional RNA-FM block to 64 PCs (Supplementary Figure S1). These were concatenated with the 15 base-model probability columns to form a 129-dimensional input vector. The meta-learner was a two-hidden-layer MLP, Layer1 (64 neurons) and Layer2 (16 neurons) trained with Adam optimization, L2 regularization and early stopping [48, 52–53]. The complete stacked architecture is summarized in Table 3; hyperparameters for all models are provided in Supplementary Table S4.

**Table 3.** Architecture of the stacked RNA sequence-based predictor. The final model combines PCA-compressed handcrafted and RNA-FM features with out-of-fold base-model probabilities.

| **Component** | **Configuration** |
| --- | --- |
| Input features | 167 handcrafted RNA features + 640 RNA-FM features |
| Base learners | 15 classifiers |
| Stacking strategy | Stratified 5-fold out-of-fold probability stacking |
| Tabular feature compression | StandardScaler + PCA, 50 components |
| RNA-FM feature compression | StandardScaler + PCA, 64 components |
| Meta-model input | 50 tabular PCs + 64 RNA-FM PCs + 15 base-model probabilities |
| Meta-model input dimension | 129 |
| Meta-classifier | Seed-bagged MLP |
| Evaluation set | Independent test set, n = 183 |

### 2.7 Model evaluation and uncertainty estimation

Model performance was assessed using accuracy, precision, recall, F1 score, Matthews correlation coefficient (MCC), area under the receiver operating characteristic curve (AUC-ROC) and area under the precision–recall curve (AUC-PR) [54–56]. Binary class labels were assigned using a fixed probability threshold of 0.5. Metric definitions are provided in Supplementary Table S5. Model stability was evaluated using stratified five-fold cross-validation within the training set. For the stacked model, cross-validation metrics were calculated from out-of-fold predictions generated for samples not used to train the corresponding models. Uncertainty in test-set performance was estimated using 2,000 bootstrap resamples of the saved test predictions [57]. Models were not retrained during bootstrapping, and 95% confidence intervals were calculated from the 2.5th and 97.5th percentiles of the bootstrap distributions.

## 3. Results

### 3.1 LLPS-associated RNAs show depletion in compositional and k-mer signature

To identify sequence features that distinguish LLPS+ from LLPS-, we compare all 167 handcrafted descriptors using two-sided Mann–Whitney U tests with Benjamini–Hochberg correction. Features with q<0.05 and ∣r_rb∣≥0.147 are considered robustly class-separating. Here, r_rb denotes the rank-biserial correlation effect size with positive values indicating higher feature values in LLPS+ RNAs. Of the 167 descriptors, 103 are significant after correction and 69 satisfy both the significance and effect-size criteria. The strongest differences involve local composition and k-mer usage with additional feature families examined in Section 3.2. Representative descriptors and their statistics are shown in Figures 2 and 3 and Supplementary Table S6.

**Figure 2.**
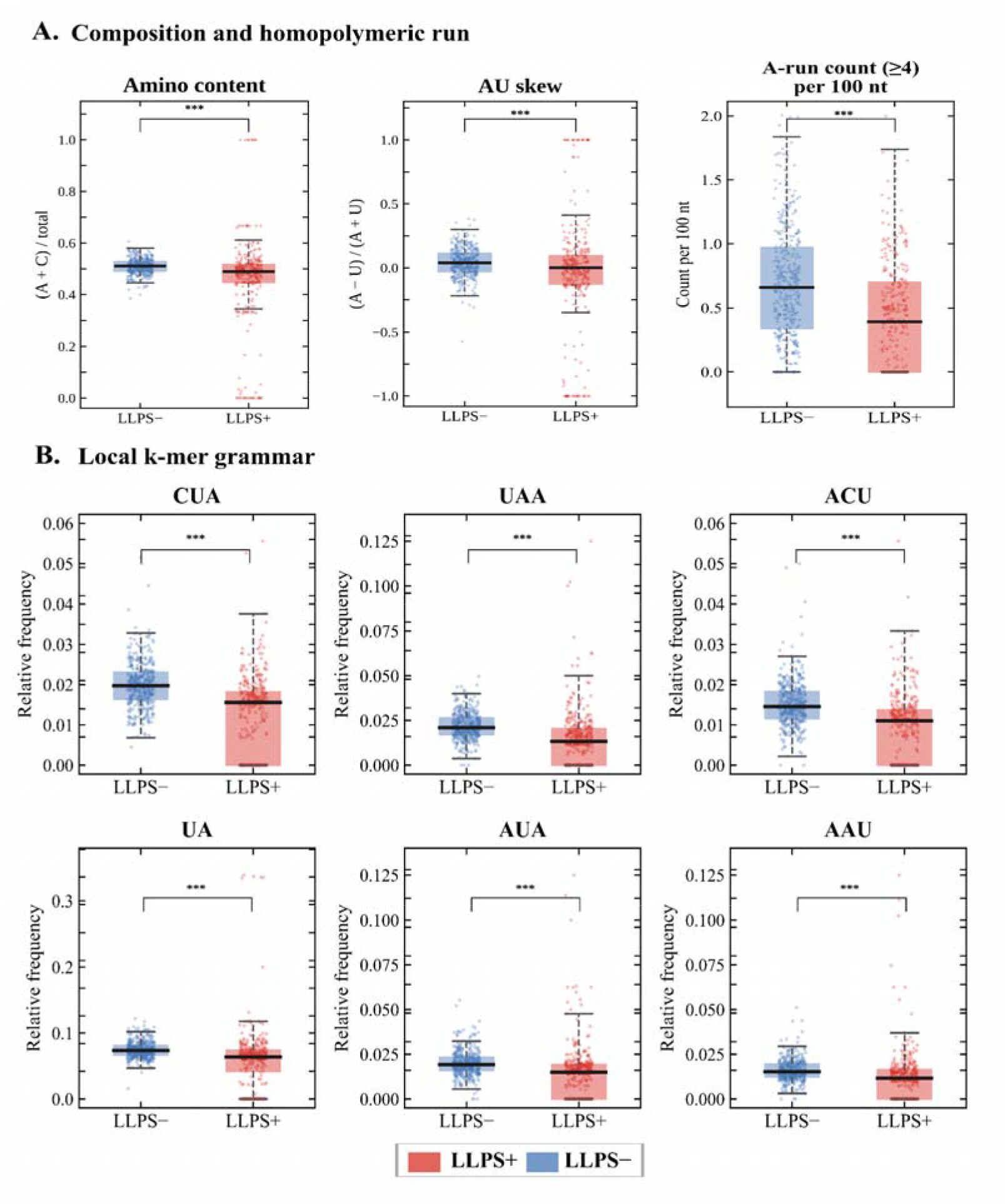
Representative composition, homopolymeric-run and local k-mer features distinguishing LLPS− and LLPS+ RNAs. Boxplots show the median and interquartile range. (A) Representative composition-related features including nucleotide composition, AU skew and A-run abundance (runs of ≥4 nt per 100 nt). (B) Representative local k-mer frequencies showing depletion in LLPS+ RNAs, including CUA, UAA, ACU, UA, AUA and AAU. Statistical comparisons are performed using two-sided Mann–Whitney U tests with Benjamini–Hochberg false-discovery-rate correction. Significance levels are indicated as follows: ns (not significant), *p < 0.05, **p < 0.01, ***p < 0.001. Full statistics are provided in Supplementary Table S3.

**Figure 3.**
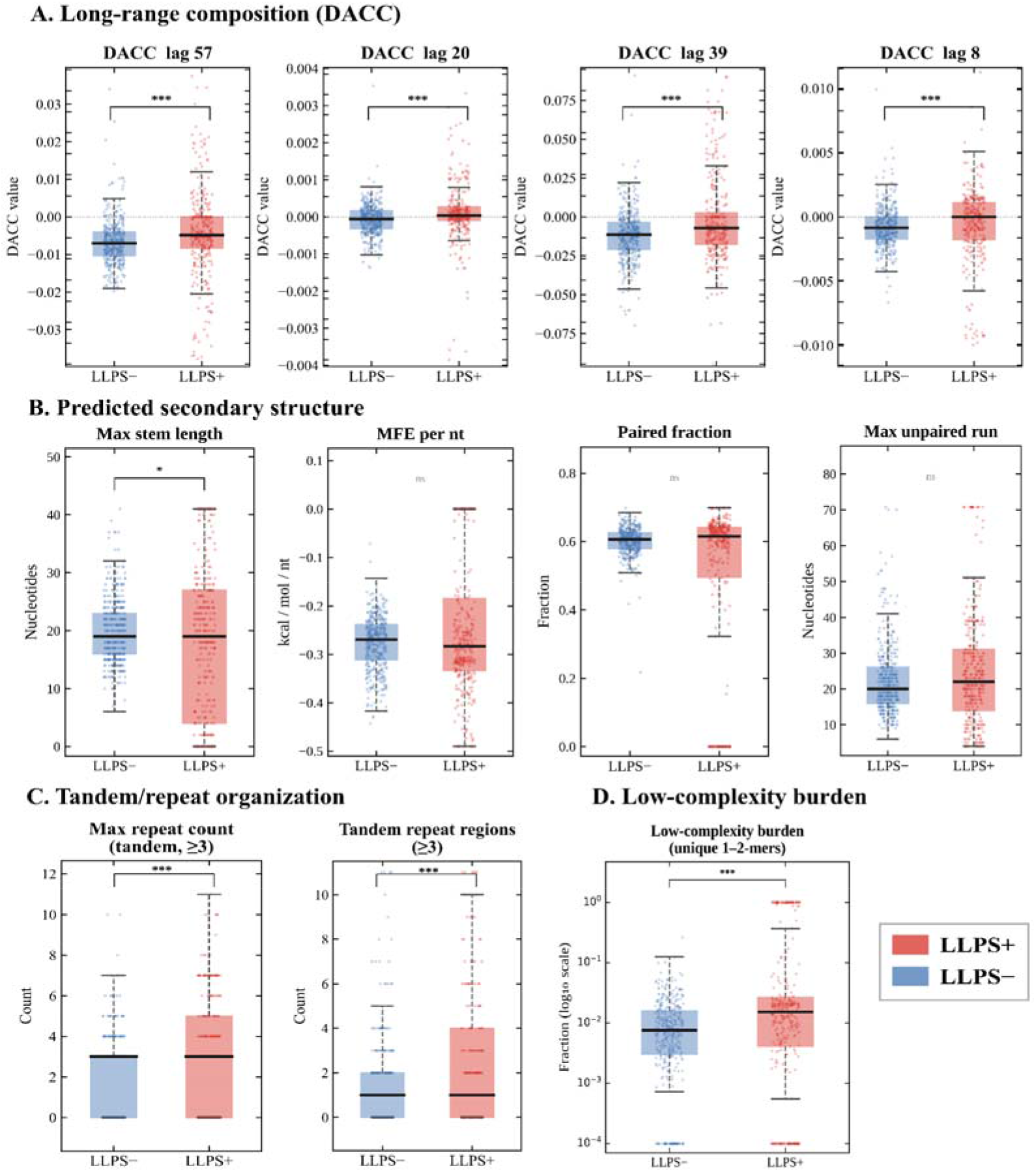
Representative long-range sequence-order, predicted secondary-structure, repeat and low-complexity descriptors comparing LLPS− and LLPS+ RNAs. Boxplots show the median and interquartile range, with individual points representing RNAs. (A) Selected DACC descriptors capturing long-range dinucleotide sequence organization at different lag distances. (B) Predicted secondary-structure descriptors derived from RNAfold including maximum stem length, minimum free energy per nucleotide, paired fraction and maximum unpaired-run length. (C) Tandem-repeat organization represented by maximum tandem-repeat count and the number of tandem-repeat regions. (D) Low-complexity burden based on unique 1–2-mer sequence content shown on a log10 scale. Statistical comparisons are performed using two-sided Mann–Whitney U tests with Benjamini–Hochberg false-discovery-rate correction. Statistical significance is evaluated using the Mann-Whitney U test; significance levels are indicated as follows: ns (not significant), *p < 0.05, **p < 0.01, ***p < 0.001. Full statistics are provided in Supplementary Tables S3.

Composition-related features show clear but generally more moderate class differences (Figure 2A). Most notably, the number of homopolymeric A-runs of length ≥4 per 100 nucleotides (A_run_count_ge4_per100nt) is significantly lower in LLPS+ RNAs, decreasing from a median of 0.659 in LLPS− RNAs to 0.393 in LLPS+ RNAs (∣r_rb∣=0.33, q=9.4×10−14). This is the only homopolymer descriptor to satisfy both thresholds and shows the largest effect size outside the composition/k-mer family. Its depletion indicates that the class difference extends beyond individual nucleotide-word frequencies to the local organization of A-rich sequence tracts.

Local k-mer usage provides the strongest and the most consistent separation between classes (Figure 2B). Among the 66 k-mer descriptors, 55 are significant after FDR correction and 47 satisfy both the significance and effect-size thresholds. The effect-size direction is negative for 63 of the 66 descriptors, indicating a broad depletion of selected sequence words in LLPS+ RNAs. The strongest individual effect is observed for CUA, which decreases from a median frequency of 0.0197 in LLPS− RNAs to 0.0156 in LLPS+ RNAs (∣r_rb∣=0.49, q=3.9×10−28). Other strongly depleted features include UAA (∣r_rb∣=0.46), ACU (∣r_rb∣=0.46), UA (∣r_rb∣=0.39) and AUA (∣r_rb∣=0.39). The depleted descriptors are predominantly A/U-containing di- and tri-nucleotides, whereas only three descriptors show higher values in LLPS+ RNAs. The dominant compositional pattern is therefore better characterized as selective depletion of A/U-rich sequence words than as enrichment of a single nucleotide class. These results identify local sequence composition as the strongest univariate source of class separation. The coordinated depletion of selected A/U-containing k-mers and the reduced abundance of homopolymeric A-runs suggest that LLPS-associated RNAs differ from background RNAs in both local sequence-word usage and short-range sequence organization.

### 3.2 LLPS-associated RNAs show altered long-range sequence order, repeat organization and predicted structural features

Beyond the dominant compositional and k-mer signal, several additional feature families show weaker differences between LLPS+ and LLPS− RNAs. These effects are generally small (∣r_rb∣<0.30) and are therefore interpreted as supporting components of a broader sequence architecture rather than as primary discriminators. Long-range sequence-order descriptors provide an additional layer of separation (Figure 3A). Thirteen DACC features satisfy both the significance and effect-size criteria, with mixed directions of change in LLPS+ RNAs. Because DACC descriptors summarize correlations among dinucleotide physicochemical properties across sequence positions, these results suggest that LLPS+ and LLPS− RNAs differ not only in the frequencies of short nucleotide words but also in how dinucleotide properties are arranged across the sequence. The modest effect sizes indicate that these differences contribute incrementally rather than defining the classes independently.

In contrast, global RNAfold-derived descriptors show limited class-wise separation (Figure 3B). Minimum free energy per nucleotide, paired fraction and maximum unpaired-run length differ little between LLPS+ and LLPS− RNAs, while maximum stem length shows only a weak, nonsignificant trend. These observations do not exclude a role for RNA structure in condensate biology, but suggest that coarse whole-sequence structural summaries may not capture local, alternative or protein-dependent structural states relevant to LLPS. Repeat-related descriptors show a more selective pattern (Figure 3C). Two tri-nucleotide tandem-repeat features are modestly elevated in LLPS+ RNAs. This contrasts with the depletion of homopolymeric A-runs described in Section 3.1, indicating that repeat-associated differences depend on repeat type rather than reflecting a uniform increase in repetitive sequence. Low-complexity burden also shows modest class separation (Figure 3D). The strongest feature in this group, low_complexity_burden_unique12, increases from a median of 0.0075 in LLPS− RNAs to 0.0152 in LLPS+ RNAs (∣r_rb∣=0.23, q=2.5×10−7). Although the absolute difference is small, the shift indicates a greater prevalence of compositionally restricted sequence regions among LLPS-associated RNAs. Such regions may increase the local recurrence of related sequence elements and thereby influence opportunities for multivalent RNA–RNA or RNA–protein interactions. Overall, these results indicate that the strongest class separation arises from local sequence composition and k-mer usage, while long-range dinucleotide organization, selected tandem-repeat features and low-complexity burden provide additional, more modest signals. Global secondary-structure summaries show comparatively little separation, supporting a view of LLPS-associated RNA sequence architecture as distributed across multiple sequence properties rather than dominated by a single feature class.

### 3.3 Complementary sequence representations and stacking improve LLPS-associated RNA classification

Having identified sequence-level differences between LLPS+ and LLPS− RNAs, we have examined whether these signals support predictive classification. Identically configured L2-regularized logistic-regression models are evaluated using four input representations: sequence length and GC fraction, handcrafted descriptors, RNA-FM embeddings and the combination of both feature blocks (Supplementary Table S7). The length and GC baseline shows limited discriminatory ability, indicating that classification is not driven primarily by these coarse properties. In contrast, handcrafted descriptors and RNA-FM embeddings are individually informative, achieving AUC-ROC values of 0.879 and 0.909, respectively. Combining the two feature blocks further improves performance to an AUC-ROC of 0.924, consistent with the representations capturing complementary aspects of RNA sequence organization. Five-fold cross-validation further shows that several individual classifiers, particularly tree-based ensemble models, achieve strong predictive performance within the training data (Table 4). The stacked Meta-NN remains competitive across evaluation metrics and shows a balanced combination of precision and recall, although it did not consistently outperform the strongest individual base learners during cross-validation. This indicates that the benefit of stacking lies less in dominating every cross-validation metric than in integrating partially distinct decision patterns from multiple classifiers together with the two sequence representations. On the independent test set, the complete stacked Meta-NN achieves an accuracy of 0.891, precision of 0.913, recall of 0.849, F1 score of 0.880, AUC-ROC of 0.935, AUC-PR of 0.944 and MCC of 0.782. These results show that LLPS-associated RNA classification benefits from combining interpretable sequence descriptors with RNA-FM representations and heterogeneous classifier outputs, while the weak length and GC baseline indicates that the predictive signal extends beyond coarse sequence composition.

**Table 4.** Five-fold cross-validation performance during model development. Base-model metrics were computed from pooled out-of-fold predictions on the training set. The stacked Meta-NN was evaluated using a secondary five-fold cross-validation procedure.

| Model | Accuracy | Precision | Recall | F1 | AUC-ROC | AUC-PR | MCC |
| --- | --- | --- | --- | --- | --- | --- | --- |
| Logistic Regression (L2) | 0.803 | 0.804 | 0.766 | 0.784 | 0.866 | 0.876 | 0.604 |
| Ridge Classifier | 0.745 | 0.730 | 0.719 | 0.725 | 0.769 | 0.770 | 0.486 |
| LDA | 0.814 | 0.863 | 0.716 | 0.783 | 0.885 | 0.881 | 0.631 |
| KNN | 0.757 | 0.825 | 0.608 | 0.700 | 0.840 | 0.828 | 0.520 |
| Gaussian NB | 0.743 | **0.919** | 0.494 | 0.643 | 0.730 | 0.763 | 0.524 |
| Random Forest | 0.828 | 0.878 | 0.734 | 0.799 | 0.898 | 0.907 | 0.659 |
| Extra Trees | **0.840** | 0.900 | 0.740 | 0.812 | **0.909** | **0.917** | **0.685** |
| Hist Gradient Boosting | 0.836 | 0.863 | 0.772 | 0.815 | 0.902 | 0.907 | 0.672 |
| Gradient Boosting | 0.827 | 0.848 | 0.766 | 0.805 | 0.898 | 0.904 | 0.652 |
| AdaBoost | 0.822 | 0.823 | **0.789** | 0.806 | 0.888 | 0.889 | 0.643 |
| LightGBM | 0.832 | 0.861 | 0.763 | 0.809 | 0.905 | 0.910 | 0.664 |
| XGBoost | 0.837 | 0.861 | 0.778 | **0.817** | 0.905 | 0.911 | 0.674 |
| CatBoost | 0.839 | 0.878 | 0.760 | 0.815 | 0.906 | 0.916 | 0.679 |
| SVM-RBF | 0.822 | 0.846 | 0.757 | 0.799 | 0.895 | 0.897 | 0.644 |
| MLP-base | 0.820 | 0.839 | 0.760 | 0.798 | 0.888 | 0.897 | 0.638 |
| **Stacked Meta-NN** | 0.835 | 0.849 | 0.787 | 0.816 | 0.902 | 0.909 | 0.668 |

### 3.4 The stacked Meta-NN achieves robust test set performance

Final model performance is evaluated on the independent test set of 183 RNAs, which is not used for feature selection, cross-validation, PCA fitting or model optimization. The stacked Meta-NN achieves the strongest overall performance among the evaluated models (Table 5). Its ROC and precision–recall curves, together with the confusion matrix at the default probability threshold of 0.5, are shown in Figure 4. The Meta-NN ranks first on six of the seven evaluation metrics, while Extra Trees achieves marginally higher precision. Its high AUC-ROC and AUC-PR indicate strong discrimination across probability thresholds, whereas the F1 score and MCC demonstrate a balanced classification of LLPS+ and LLPS− RNAs. Overall, the stacked framework provides the best trade-off between ranking ability, sensitivity and classification accuracy. A metric-level comparison of all base learners and the Meta-NN is shown in Supplementary Figure S2. To quantify uncertainty in the independent test set performance estimates, we perform bootstrap resampling of the test-set predictions. Across 2,000 bootstrap resamples, the stacked Meta-NN retains robust performance (Supplementary Table S8). These confidence intervals indicate that the observed test-set performance is stable under resampling. Bootstrap confidence intervals for AUC-ROC across models are visualized in Supplementary Figure S3.

**Figure 4.**
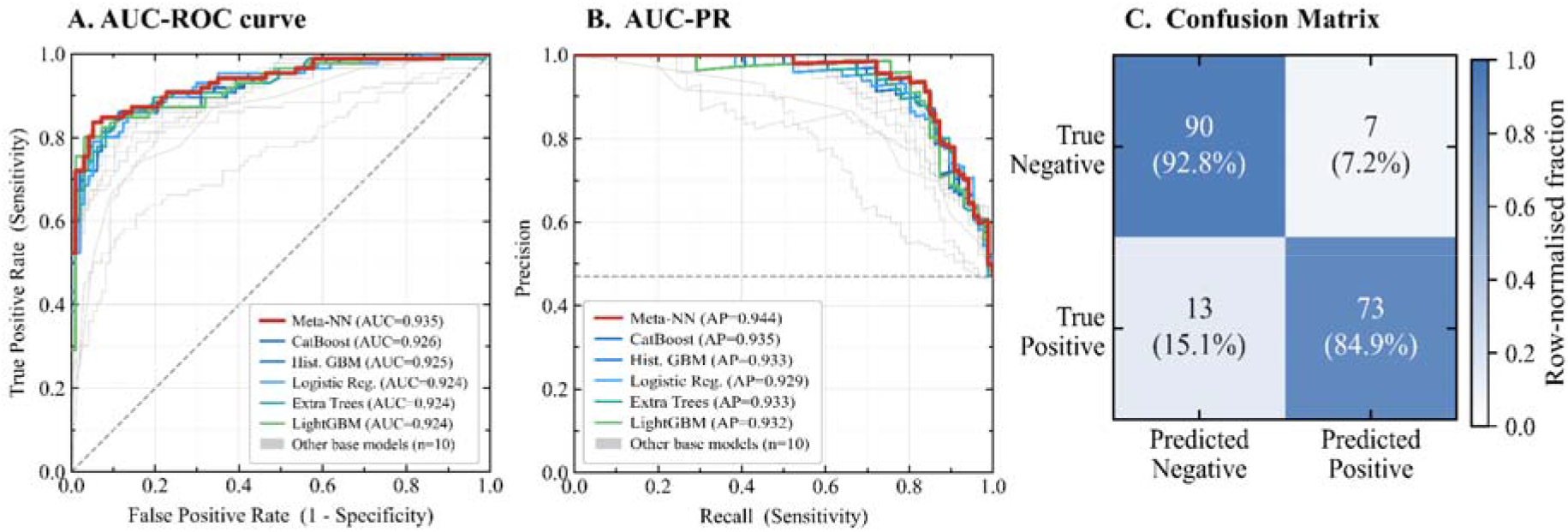
Independent test performance of the stacked Meta-NN. Performance of the final stacked Meta-NN compared with representative base classifiers on the independent held-out test set. (A) ROC curves showing true positive rate against false positive rate. (B) Precision–recall curves showing precision across recall values. (C) Row-normalized confusion matrix for the stacked Meta-NN at the default probability threshold of 0.5. Grey curves in panels A and B represent the remaining base models.

**Table 5.** Independent test-set performance of all base learners and the final stacked predictor. Metrics were computed on the independent test set of 183 RNAs. Bold values indicate the best value for each metric.

| **Model** | **Accuracy** | **Precision** | **Recall** | **F1** | **AUC-ROC** | **AUC-PR** | **MCC** |
| --- | --- | --- | --- | --- | --- | --- | --- |
| Logistic Regression (L2) | 0.853 | 0.839 | 0.849 | 0.844 | 0.924 | 0.930 | 0.704 |
| Ridge Classifier | 0.798 | 0.803 | 0.756 | 0.778 | 0.825 | 0.834 | 0.594 |
| LDA | 0.842 | 0.890 | 0.756 | 0.818 | 0.919 | 0.921 | 0.686 |
| KNN | 0.792 | 0.864 | 0.663 | 0.750 | 0.854 | 0.838 | 0.592 |
| Gaussian NB | 0.721 | 0.907 | 0.453 | 0.605 | 0.760 | 0.781 | 0.485 |
| Random Forest | 0.869 | 0.908 | 0.802 | 0.852 | 0.920 | 0.928 | 0.740 |
| Extra Trees | 0.863 | **0.918** | 0.779 | 0.843 | 0.924 | 0.933 | 0.731 |
| Hist Gradient Boosting | 0.869 | 0.888 | 0.826 | 0.855 | 0.925 | 0.933 | 0.737 |
| Gradient Boosting | 0.863 | 0.868 | 0.837 | 0.852 | 0.917 | 0.931 | 0.726 |
| AdaBoost | 0.831 | 0.831 | 0.802 | 0.817 | 0.898 | 0.913 | 0.660 |
| LightGBM | 0.869 | 0.888 | 0.826 | 0.855 | 0.924 | 0.932 | 0.737 |
| XGBoost | 0.863 | 0.877 | 0.826 | 0.850 | 0.919 | 0.932 | 0.726 |
| CatBoost | 0.863 | 0.877 | 0.826 | 0.850 | 0.926 | 0.935 | 0.726 |
| SVM-RBF | 0.847 | 0.863 | 0.802 | 0.831 | 0.919 | 0.923 | 0.693 |
| MLP-base | 0.820 | 0.835 | 0.767 | 0.800 | 0.901 | 0.910 | 0.638 |
| **Stacked Meta-NN** | **0.891** | 0.913 | **0.849** | **0.880** | **0.935** | **0.944** | **0.782** |

### 3.5 Exploratory ablation analysis of stacked-model components

To examine how individual components affected predictive performance, we performed an exploratory ablation analysis of eight architectural configurations (Figure 5; Supplementary Table S9). The ablation therefore assesses the information contributed by the handcrafted components, RNA-FM components and base-model probabilities at the meta-learning stage. Config A, represents the strongest non-stacked model, achieves an AUC-ROC of 0.919, an F1 score of 0.850 and an MCC of 0.726. Config B uses only the 15 base-model probability outputs as meta-learner inputs and improves these values to 0.932, 0.874 and 0.770, respectively. This improvement is consistent with the heterogeneous classifiers capturing complementary prediction patterns. Adding either feature block separately does not produce further gains in threshold-dependent performance. Config C combines handcrafted PCA components with the base-model probabilities and achieves an F1 score of 0.869, whereas Config D combines RNA-FM PCA components with the base-model probabilities and reaches an F1 score of 0.868. Nevertheless, Config D produces the highest AUC-ROC among all configurations (0.937), indicating strong ranking performance. These findings suggest that much of the information from either feature block already represented in the base-model probability outputs when each block is added alone. Config E, which uses 64-component PCA representation together with the base-model probabilities, performs similarly to Config B but does not match the final block-wise architecture.

**Figure 5.**
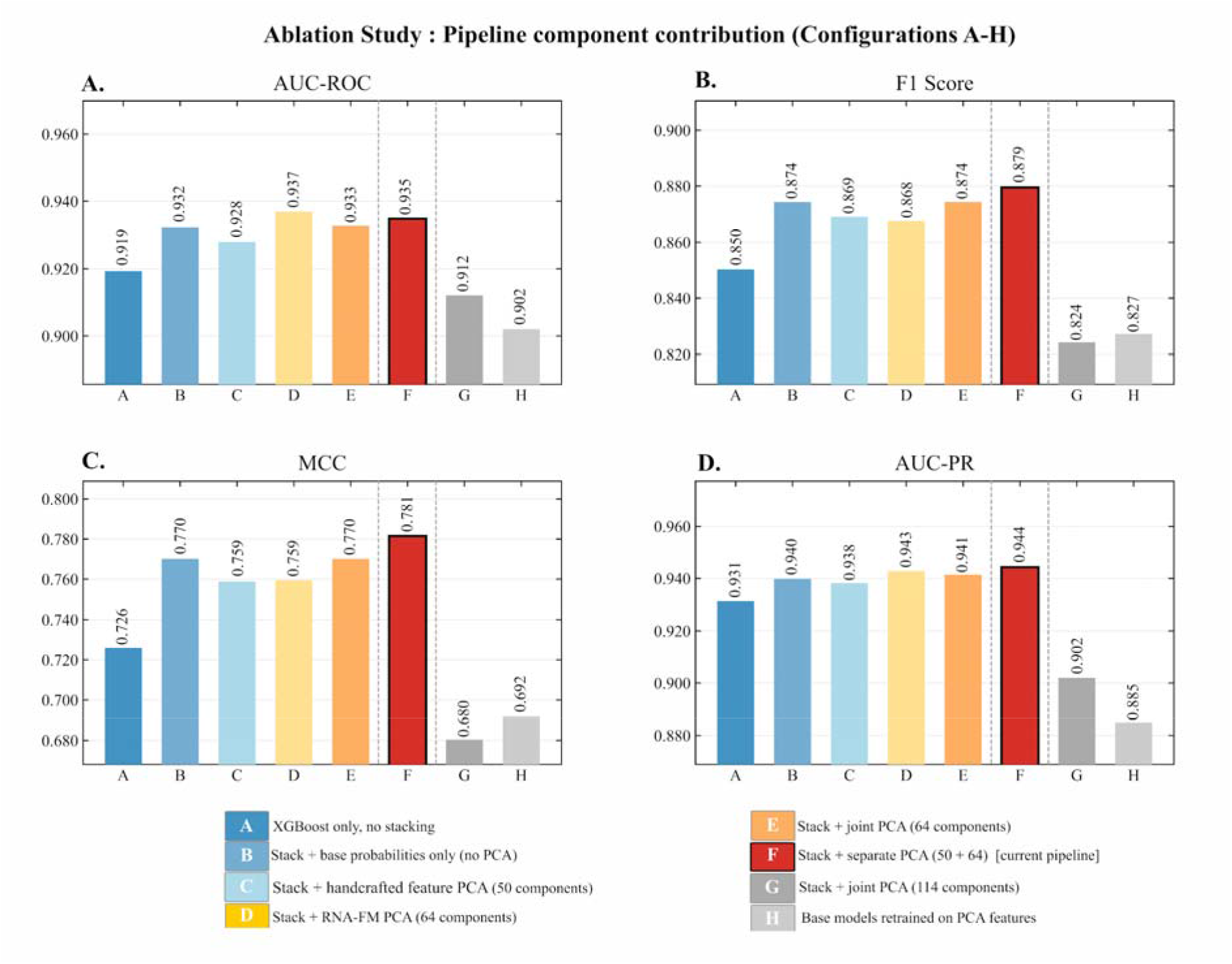
Ablation analysis of pipeline components. Performance across ablation configurations A–H. Bars show performance for (A) AUC-ROC, (B) F1 score, (C) MCC and (D) AUC-PR. Configuration F represents the final model, combining base-model probabilities with separate PCA-compressed handcrafted and RNA-FM feature blocks. The separate-PCA stack showed the strongest overall balance across threshold-dependent and ranking metrics.

The final architecture, Config F, uses separate PCA compression of handcrafted and RNA-FM feature blocks and concatenated both with the 15 base-model probability features. This configuration achieves the best performance, with the highest accuracy (0.891), F1 (0.880), AUC-PR (0.944) and MCC (0.782). Although Config D shows a slightly higher AUC-ROC, Config F provides the best overall balance across threshold-dependent and ranking metrics. In contrast, Config G, which uses joint PCA with 114 components to match the dimensionality of the separate PCA blocks, performs worse, suggesting that simply increasing joint PCA dimensionality does not recover the benefit of preserving feature-block-specific structure. Finally, Config H, where base models are trained directly on PCA-compressed features, underperform the final separate-PCA stack, indicating that PCA compression is the most useful at the meta-model integration stage rather than as a replacement for the original feature space in base-model training. These ablations show that the final stacked predictor benefits from three complementary information sources: base-model probability patterns, PCA-compressed handcrafted descriptors and PCA-compressed RNA-FM embeddings.

### 3.6 SHAP analysis identifies complementary handcrafted and RNA-FM predictive signals

To interpret the features contributing to LLPS-associated RNA classification, we perform SHAP analysis [58–59] using a LightGBM model trained on the complete 807-feature representation. Features are ranked by mean absolute SHAP value, which measures their average contribution to model predictions across RNAs without directly implying class-wise enrichment or depletion. The highest-ranked features include both handcrafted descriptors and RNA-FM embedding dimensions, indicating that the interpretation model draws on complementary explicit and latent sequence information (Figure 6A; Supplementary Table S10). Among the handcrafted descriptors, the highest-ranked feature is linguistic_complexity_k1_5, followed by selected k-mer features including CUU, CUA, AUU, UAU_density and UAA, as well as the DACC and repeat-related descriptors dacc_21, dacc_24 and U_run_count_ge4. These feature categories are consistent with the statistical analysis, which identifies altered k-mer usage, sequence complexity, repeat organization and dinucleotide-order patterns as signatures distinguishing LLPS+ from LLPS− RNAs.

**Figure 6.**
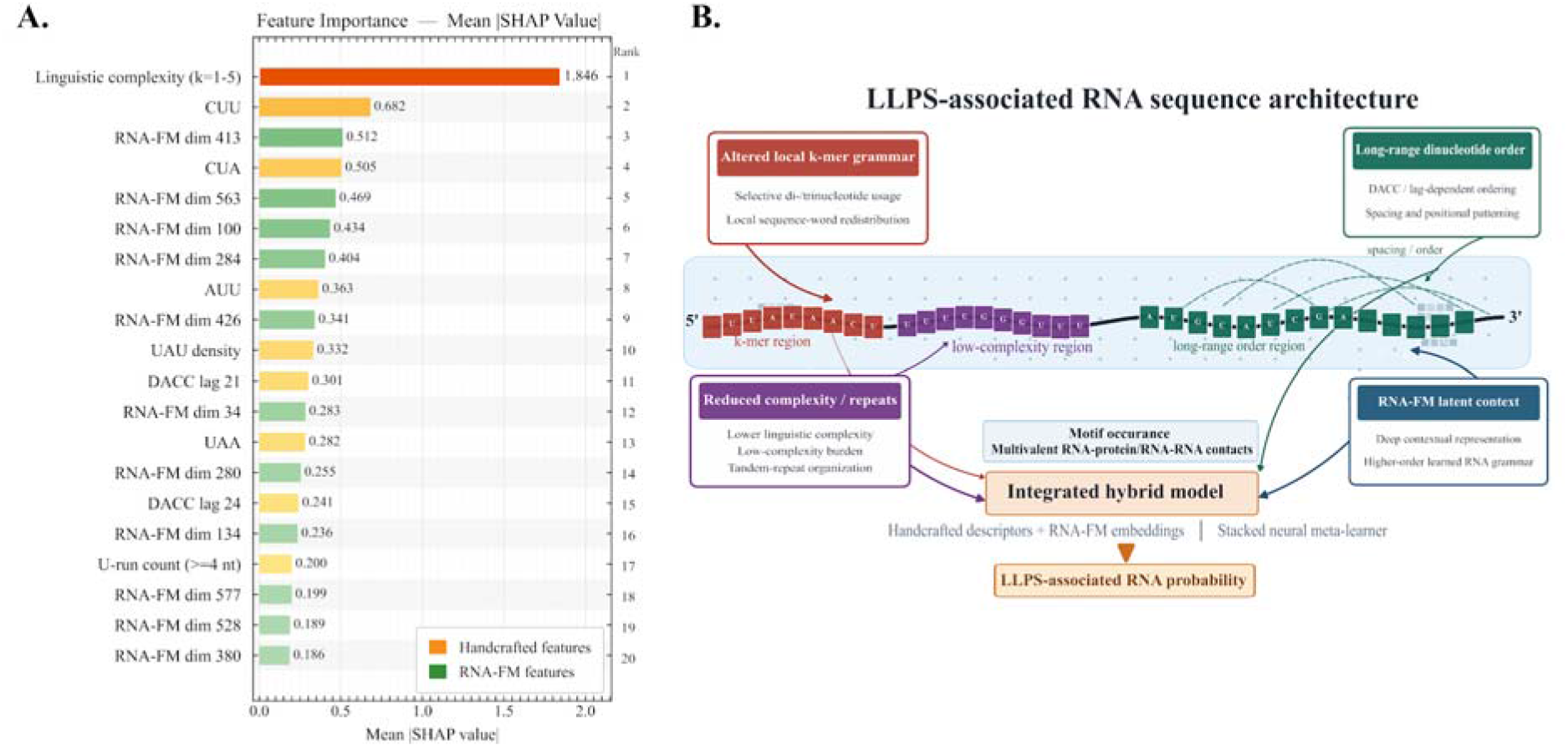
SHAP-based interpretation and proposed sequence architecture of LLPS-associated RNAs. (A) Top model-contributing features ranked by mean absolute SHAP value from a LightGBM model trained on the full 807-feature representation. Colours indicate handcrafted descriptors or RNA-FM embedding dimensions. (B) Proposed interpretation of the major feature categories supporting LLPS-associated RNA prediction including altered local k-mer grammar, reduced sequence complexity and repeat organization, long-range dinucleotide order and RNA-FM latent sequence context.

RNA-FM dimensions 413, 563, 100, 284, 426, 34, 280, 134, 577, 528 and 380 also appear prominently among the top-ranked features. These dimensions are not assigned direct biological meanings because they represent distributed latent coordinates learned by the pretrained RNA language model. Nevertheless, their prominence suggests that RNA-FM captures contextual sequence information that complements the handcrafted descriptors. Mean absolute SHAP values should be interpreted as measures of predictive contribution in a multivariate model rather than as evidence that individual features are independently enriched or depleted in LLPS+ RNAs. Direct class-wise comparisons are reported in Supplementary Table S3, while representative plotted features are summarized in Supplementary Table S6. Overall, the SHAP results support the combined use of biologically interpretable sequence descriptors and latent RNA-FM representations. The major feature categories are summarized schematically in Figure 6B.

### 3.7 Representative biological case studies of model predictions

To examine model behaviour at the level of individual RNAs, we consider representative correctly classified and misclassified examples from the independent test set. Among correctly classified LLPS-associated RNAs, MALAT1 is predicted as LLPS+ with a Meta-NN probability of 0.957. Its sequence profile does not uniformly follow the dominant depletion of A/U-containing k-mers observed across the LLPS+ class (Section 3.1); instead, its A-run abundance, linguistic complexity and selected DACC descriptors are more consistent with the broader LLPS-associated profile. TMPO-AS1 is also correctly classified as LLPS+ (probability = 0.710) and shows several features consistent with the dominant class-level tendencies including depletion of selected A/U-containing k-mers, reduced A-run abundance and increased low-complexity burden (Sections 3.1–2.2). Together, these examples illustrate that LLPS-associated predictions can arise from different combinations of local and broader sequence properties rather than from a single universal feature pattern.

Among misclassified RNAs, HOTAIRM1 is labelled LLPS+ but predicted as LLPS− with a low Meta-NN probability and limited support across the base classifiers. This suggests that HOTAIRM1 does not strongly match the dominant sequence signature learned by the model, including altered k-mer usage, sequence-complexity features and repeat-associated organization. HOTAIRM1 functions in DNA double-strand break repair through association with Y14/RBM8A and Ku70/80-containing repair factors, while Y14–RNA co-phase separation is also implicated in DNA-damage repair [60–62]. HOTAIRM1 may therefore represent an LLPS-associated RNA whose condensate participation depends more strongly on protein-mediated recruitment, localization or other cellular context not fully captured by primary sequence alone. As a representative false-positive case, the background-labelled mouse lncRNA Gm56852 (ENSMUST00000241776), annotated in Ensembl [21], is predicted as LLPS+ with broad ensemble support. Manual inspection does not identify an exact reviewed LLPS-positive match by sequence, transcript ID, gene ID or same-organism annotation. The transcript is therefore retained in the LLPS− background class under the benchmark definition (Section 2.1). Nevertheless, its sequence contains compositional and low-complexity properties resembling those observed among LLPS-associated RNAs. Because the LLPS− class represents LLPS-unannotated background rather than experimentally verified non-LLPS RNAs, this prediction may represent either a genuine model error or an unannotated RNA carrying LLPS-like sequence properties. These cases illustrate both the heterogeneity and the limitations of sequence-based LLPS-RNA prediction. Correctly classified LLPS-associated RNAs can exhibit different combinations of local and broader sequence properties, whereas false negatives may reflect condensate association that depends more strongly on protein recruitment, localization or condition-specific mechanisms. Apparent false positives may similarly represent either genuine model errors or LLPS-like sequence properties in currently background RNAs.

## 4. Discussion

### 4.1 LLPS-associated RNAs exhibit a distributed sequence architecture captured by complementary representations

Computational studies of LLPS have largely focused on protein sequence features or on defined protein–RNA systems under specific experimental conditions. The present study addresses a related but distinct question: whether experimentally supported LLPS-associated RNAs exhibit reproducible primary-sequence signatures and whether these signatures can support classification from RNA sequence alone. Our findings suggest that the relevant signal is distributed across several layers of sequence organization rather than concentrated in a single feature type. LLPS-associated RNAs differ from background RNAs primarily in local k-mer usage, low-complexity burden and long-range dinucleotide autocorrelation (Figure 2-3; Supplementary Table S11). In contrast, a baseline restricted to sequence length and GC fraction provides little predictive discrimination, while the evaluated global RNAfold-derived summaries show limited class separation. The most prominent pattern involves depletion of several A/U-containing di- and tri-nucleotides, accompanied by smaller differences in poly(A) runs, localized sequence complexity and broader positional organization. These feature families have plausible biological relevance. Local k-mer usage may influence RNA–protein recognition, base-pairing potential and short structural motifs [5, 7–8]. While low-complexity and repeat-associated regions may present recurring interaction elements that facilitate multivalent RNA–protein or RNA–RNA contacts [5, 7–8, 10]. DACC descriptors provide a complementary view by capturing how dinucleotide properties are arranged across the transcript rather than simply how frequently they occur. The identified signatures therefore point toward an RNA architecture involving both the identity of local sequence elements and their organization over longer sequence scales.

Importantly, these associations should not be interpreted as evidence for a universal sequence code that is sufficient to drive LLPS. Rather, they describe statistical tendencies shared across a heterogeneous collection of experimentally supported LLPS-associated RNAs. Such tendencies may influence the capacity of an RNA to present multiple interaction sites or organize multivalent RNA–protein and RNA–RNA contacts, while the actual condensate behaviour remains dependent on protein partners, concentration, localization and cellular context. The distributed nature of this signal also provides a rationale for combining handcrafted descriptors with RNA-FM embeddings. Handcrafted features encode explicit and biologically interpretable sequence properties, whereas RNA-FM provides distributed contextual representations learned from large-scale RNA sequence data [19–20]. Their combination outperforms either feature block alone (Section 3.3), consistent with partial complementarity between predefined descriptors and learned sequence context. The ablation analysis further supports this interpretation (Figure 5; Supplementary Table S9). Configurations combining base-model probabilities with either handcrafted PCA or RNA-FM PCA alone capture part of the predictive signal, whereas Config F, which integrates both separately compressed feature blocks with the base-model probabilities, achieves the strongest overall performance. Its advantage over the base-probability-only, single-block and joint-PCA configurations suggests that the two representations contribute most effectively when integrated while retaining their distinct feature-space structures. Separate PCA compression may help preserve representation-specific information, although the present analysis does not establish the mechanism underlying this advantage. Overall, these results support the complementary use of interpretable sequence descriptors and latent RNA-FM representations for capturing the distributed sequence signal associated with LLPS.

### 4.2 Sequence organization provides mechanistic context for multivalent RNA–protein assembly

The sequence patterns identified in Sections 3.1–2.2 are broadly consistent with experimentally characterized RNA–protein condensate systems in which the number, recurrence and organization of RNA-binding elements influence multivalent interactions. These systems provide mechanistic context for the present findings without establishing that the specific sequence features identified here directly determine condensate formation. For example, Elguindy and Mendell [63] show that the lncRNA NORAD contains repeated PUMILIO-response elements that create a multivalent platform for recruitment of PUM1 and PUM2 and promote formation of phase-separated NORAD–PUMILIO bodies. Similarly, TDP-43 phase-separation behaviour is sensitive to the sequence and organization of its RNA-binding sites. Grese et al. [64] show that GU-rich RNAs containing multiple adjacent TDP-43-binding sites promote TDP-43 phase separation with condensation increasing as the number of binding sites increases. Extending this principle to longer cellular RNAs, Hallegger et al. [65] identify condensation-dependent TDP-43 assembly across extended RNA regions containing dispersed clusters of UG-rich recognition motifs, termed “binding-region condensates.”

At the structural level, the tandem RNA-recognition motifs of TDP-43 engage UG-rich RNA as illustrated in the representative TDP-43–RNA complex (Figure 7A, PDB id: 4BS2) [66]. MALAT1 provides a particularly relevant example because it contains several TDP-43-binding regions with distinct condensation dependencies [65] and also identified as a correctly classified LLPS-associated RNA in the present study (Section 3.7). The schematic in Figure 7B illustrates the dispersed organization of TDP-43-binding regions along MALAT1 and their potential contribution to multivalent TDP-43 recruitment and condensation-dependent assembly. Importantly, the sequence descriptors used here do not identify TDP-43-binding sites directly. Rather, features such as local k-mer usage, sequence complexity and longer-range sequence-order descriptors capture broader aspects of RNA organization that influence the distribution or recurrence of potential interaction regions. A complementary example is provided by TMPO-AS1, which is experimentally linked to FUS–p300 condensate formation. Luo et al. [67] show that TMPO-AS1 interacts with FUS and recruits p300 to the TMPO promoter, while depletion of TMPO-AS1 reduces colocalized nuclear FUS–p300 puncta. This experimentally characterized interaction is summarized schematically in Figure 7C, where representative structures of the FUS RRM and p300 are included for molecular context [68–69]. Together with the MALAT1–TDP-43 system, this example illustrates that condensate-associated RNAs can participate in distinct protein-dependent assemblies while exhibiting different combinations of sequence properties. These experimentally characterized systems support a model in which RNA sequence organization may influence condensate participation by modulating the number, recurrence and spatial arrangement of potential RNA–protein interaction opportunities. Such effects could contribute to multivalent recruitment without requiring a single universal LLPS motif.

**Figure 7.**
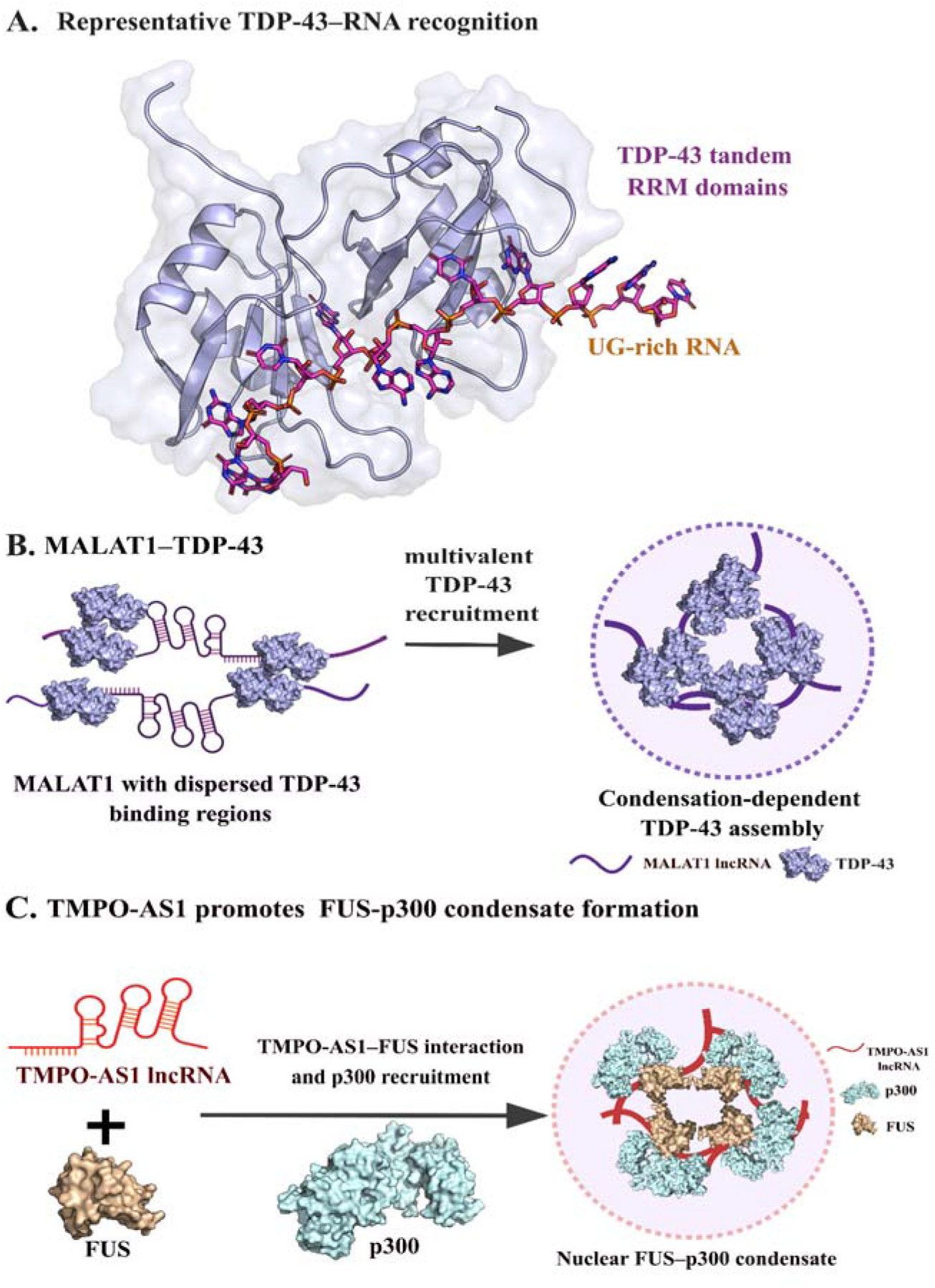
Experimentally characterized protein-RNA systems provide mechanistic context for LLPS-associated RNA sequence organization. (A) Representative structure of TDP-43–RNA recognition (PDB id: 4BS2), showing the tandem RNA-recognition motifs of TDP-43 bound to UG-rich RNA. The structure illustrates a molecular mode of sequence-specific TDP-43–RNA recognition and is not MALAT1-specific. (B) Schematic representation of MALAT1 containing dispersed TDP-43-binding regions. Experimental studies have shown that the number and organization of TDP-43 RNA-binding sites can influence multivalent recruitment and condensation-dependent TDP-43 assembly [64,65]. The TDP-43 surface structures are shown for representative molecular context and do not depict a resolved MALAT1–TDP-43 complex. (C) Schematic representation of TMPO-AS1-mediated FUS–p300 condensate formation. TMPO-AS1 interacts with FUS and promotes recruitment of p300, supporting formation of nuclear FUS–p300 condensates [66]. Surface representations of FUS and p300 are included for structural context.

### 4.3 Stacking improves prediction by combining complementary decision rules

The performance of the stacked model suggests that LLPS-associated RNA prediction benefits from combining multiple classifiers rather than relying on a single decision rule. This is consistent with the heterogeneous nature of the sequence signal identified above. The sequence signal spans local k-mer usage, low-complexity burden, repeat organization, long-range dinucleotide autocorrelation and RNA-FM-derived sequence context. Different classifier families may emphasize different aspects of this signal. Tree-based models are well suited to nonlinear and threshold-like effects, such as changes in k-mer abundance, low-complexity burden or repeat-associated descriptors. Linear and kernel-based models capture broader global separation in the feature space. While probabilistic and distance-based models may contribute additional information from alternative assumptions about class structure. Thus, different base learners may emphasize different aspects of LLPS-associated RNA sequence organization. This complementarity is reflected most clearly in the independent test set performance. As shown in Table 5, the stacked meta-learner outperformed all individual base models across the main classification metrics, achieving the highest AUC-ROC, AUC-PR, F1 score, MCC and accuracy. This pattern suggests that stacking does not simply optimize a single metric or exploit a favorable test split. Rather, it provides a robust integration of partially complementary classifier outputs. Correlations among base-model probability outputs further show shared but non-identical prediction patterns among classifiers (Supplementary Figure S4). The gain from stacking should also be viewed in light of the biological complexity of the prediction task. LLPS-associated RNA identity is unlikely to be captured by a single feature class or decision boundary, but instead appears to reflect a distributed combination of local and broader sequence properties. A stacked architecture is therefore well suited to integrate partially distinct prediction patterns from models with different inductive biases.

### 4.4 SHAP analysis links model predictions to biologically interpretable sequence features

The SHAP analysis provides a bridge between predictive modelling and biological interpretation (Figure 6A-B). The top-ranked handcrafted features include linguistic complexity, selected k-mers, DACC descriptors and repeat-related features. These categories are consistent with the independent statistical feature analysis. The analysis identifies altered k-mer grammar, reduced sequence complexity, tandem-repeat organization and long-range sequence-order effects as major signatures of LLPS-associated RNAs. This agreement increases confidence that the model is not relying on arbitrary or uninterpretable signals alone. Instead, many predictive features are aligned with biologically interpretable RNA sequence properties. At the same time, RNA-FM dimensions are also prominent among top SHAP-ranked features, supporting the view that the model used both explicit and latent sequence signals. Thus, the model performance does not arise only from black-box embeddings or only from handcrafted features, rather, from complementary information contributed by both representation types. However, SHAP results must be interpreted carefully. Mean absolute SHAP values indicate features that contribute strongly to model predictions in a multivariate setting. They do not necessarily mean that the same features are independently enriched or depleted in LLPS+ RNAs in a univariate sense. Therefore, SHAP analysis should be interpreted together with the statistical feature tables.

### 4.5 Biological implications for RNA-mediated condensate formation

The present framework supports a view in which LLPS-associated RNAs carry distributed sequence features rather than a single universal motif. Altered k-mer usage, repeat organization, sequence complexity and long-range dinucleotide pattern influence RNA–protein and RNA–RNA interactions by altering the distribution and accessibility of potential binding sites [5, 7–8, 10]. The representative error cases further suggest that LLPS-associated RNAs may arise through different molecular routes. Some may exhibit intrinsic sequence features associated with multivalent interactions, whereas others may depend more strongly on protein-mediated recruitment, localization, transcriptional state, stress responses or RNA modifications [9, 60–61]. This diversity is consistent with the improved performance obtained by combining handcrafted descriptors, RNA-FM embeddings and stacked classifier outputs. An important limitation is the relatively small and still developing set of experimentally characterized LLPS-associated RNAs. Curated RNA datasets remain less extensive than corresponding protein LLPS resources, which limits the range of mechanisms represented during model development. The predictor is therefore best viewed as a candidate-prioritization framework rather than a definitive annotation tool. As additional experimentally supported RNAs become available, future models could be evaluated on larger and externally derived benchmarks and extended with complementary information such as RNA accessibility, RNA–protein interaction maps, RBP motifs, RNA modifications, localization and expression. Experimental perturbation of selected sequence features and high-confidence predictions could further clarify their relationships with condensate enrichment, RNA localization and RNA–protein co-partitioning.

### 4.6 Conclusion

We investigate the molecular signatures that drives to distinguish LLPS-associated RNAs from LLPS-unannotated background RNAs. By combining biologically interpretable handcrafted descriptors with contextual RNA-FM language-model embeddings, we developed a hybrid machine-learning framework that integrates explicit RNA sequence architecture with latent sequence representations learned from large-scale RNA data. The stacked Meta-NN achieves strong independent test-set performance (AUC-ROC = 0.935, F1 = 0.880 and MCC = 0.782) while retaining interpretability through feature statistics, ablation analysis and SHAP-based model interpretation. Beyond prediction, the results reveal a coordinated RNA sequence architecture associated with LLPS. This architecture spans altered local k-mer grammar, increased low-complexity burden, repeat-associated organization and long-range dinucleotide patterning, together with broader RNA-FM-derived sequence context. These features suggest that LLPS-associated RNAs may carry distributed sequence-level signatures related to multivalency, motif recurrence and sequence-order effects, rather than being defined by a single motif or global compositional bias. More broadly, this framework provides a basis for prioritizing candidate LLPS-associated RNAs and for generating experimentally testable hypotheses about the sequence features that contribute to RNA-mediated condensate biology.

## Acknowledgments

SC thanks Indian Institute of Technology Kharagpur for fellowship and infrastructure support. RPB is thankful for funding from the Department of Biotechnology (DBT), Govt. of India (grant no. BT/PR40175/BTIS/137/41/2022) for the Bioinformatics Centre.

## Author Contributions

RPB and SC conceived the idea and designed the experiments. SC performed the experiments and analyses. All the authors contributed to the manuscript writing.

## Competing Interest Statement

The authors declare no competing interests.

## Data Availability

The datasets and analysis code supporting this study are publicly available in the GitHub repository: https://github.com/SonaliChatterjee-10/RNAPhase-ML

## Notes

### Competing Interest Statement

The authors have declared no competing interest.

