## Supplementary Information for "Sequence architecture of RNA reveals key molecular signatures driving RNA phase separation"

for

**Supplementary Figure S1.**

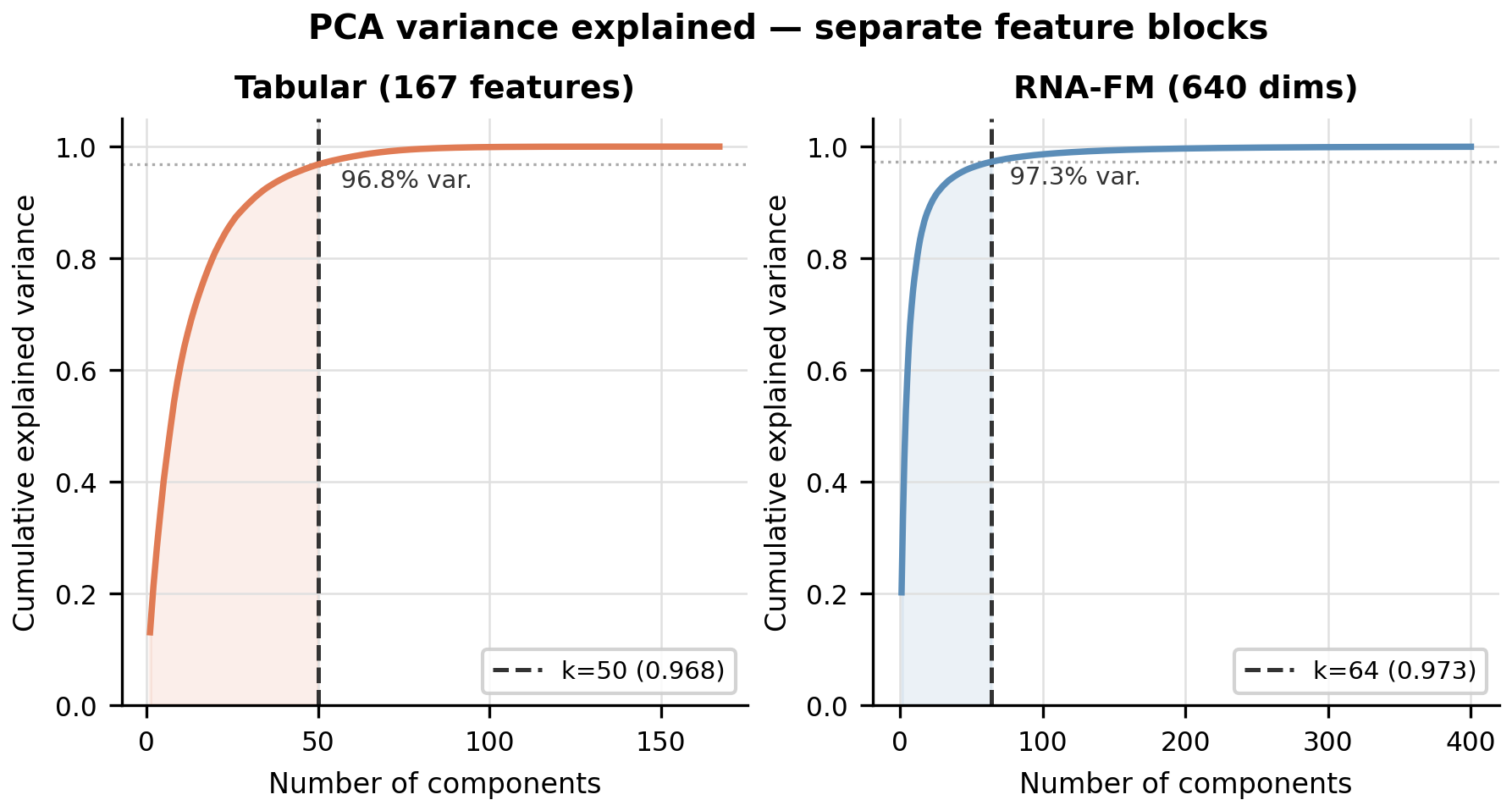

Supplementary Figure S1. PCA variance explained for separate feature blocks. Cumulative explained variance for the handcrafted descriptor block and RNA-FM embedding block. The final Meta-NN used 50 handcrafted principal components retaining 96.8% variance and 64 RNA-FM principal components retaining 97.3% variance.

**Supplementary Figure S2**

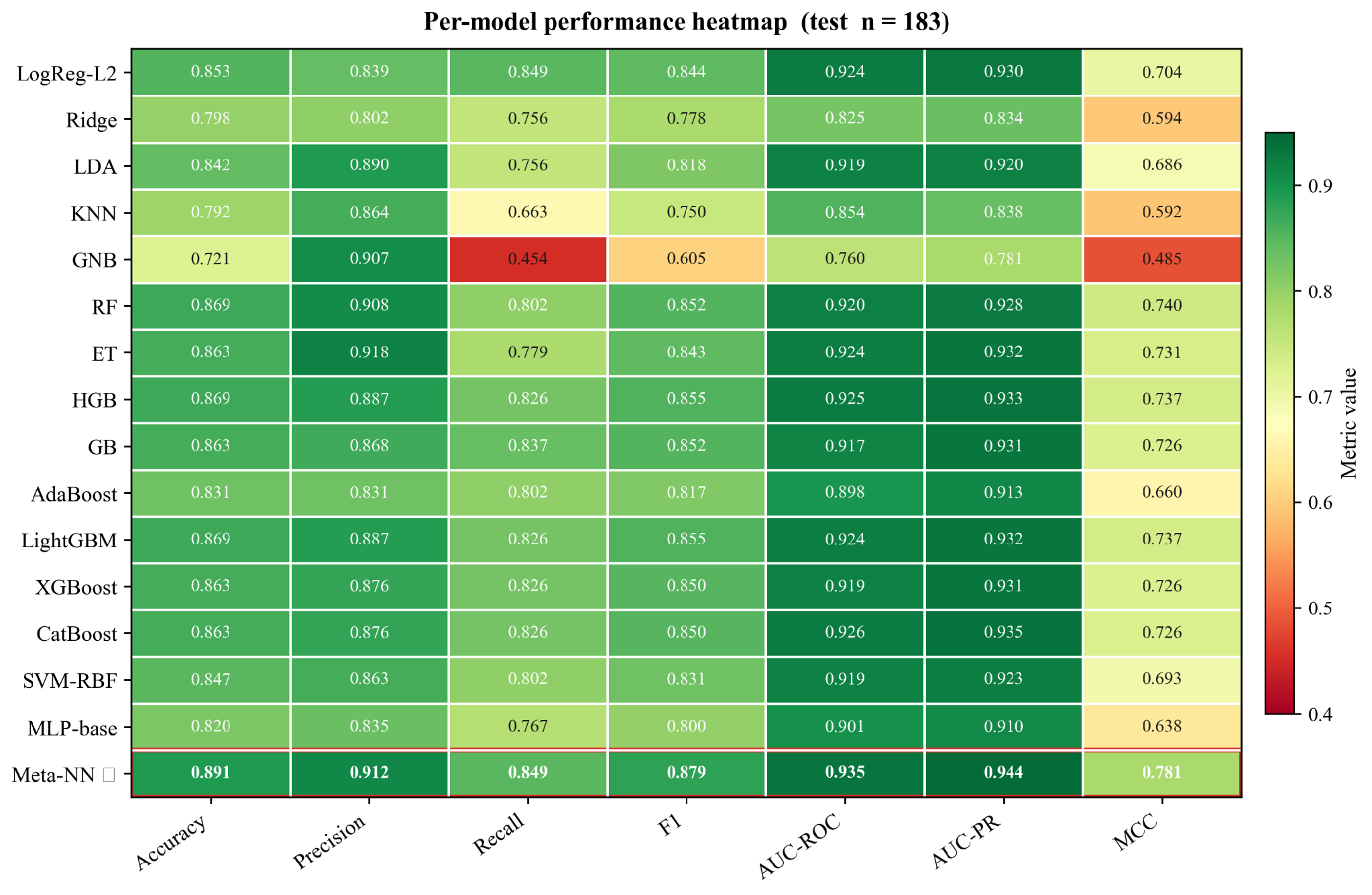

Supplementary Figure S2. Independent test set performance heatmap across all models**.**
Metric-level comparison of all base learners and the stacked Meta-NN on the independent test set. The stacked Meta-NN showed the strongest overall balance across accuracy, F1 score, AUC-ROC, AUC-PR and MCC

**Supplementary Figure S3.**

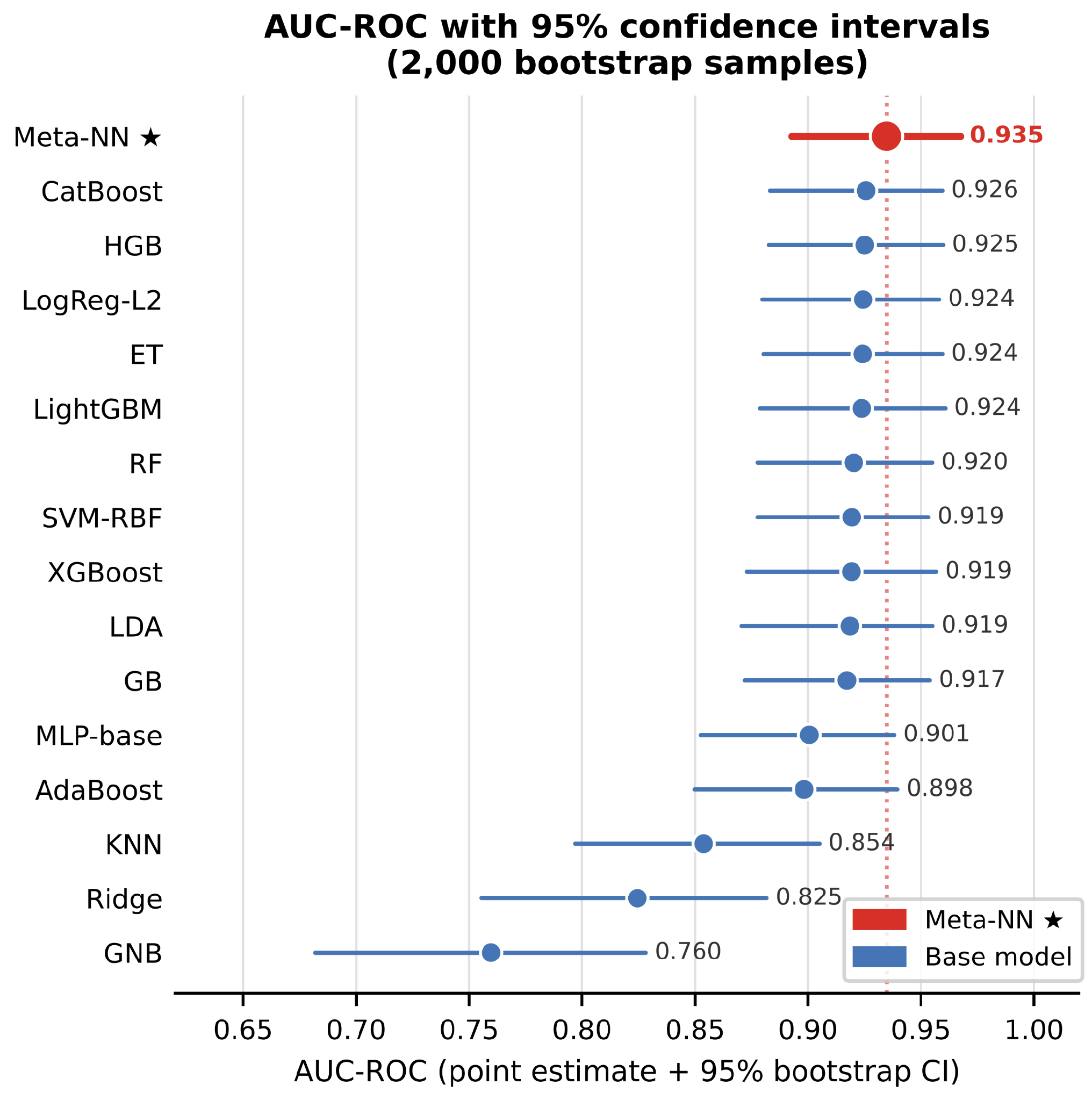

Supplementary Figure S3. AUC-ROC bootstrap confidence intervals across models. Point estimates and 95% bootstrap confidence intervals for AUC-ROC values across the stacked Meta-NN and base models. Confidence intervals were estimated using 2,000 bootstrap resamples of independent test-set predictions.

**Supplementary Figure S4.**

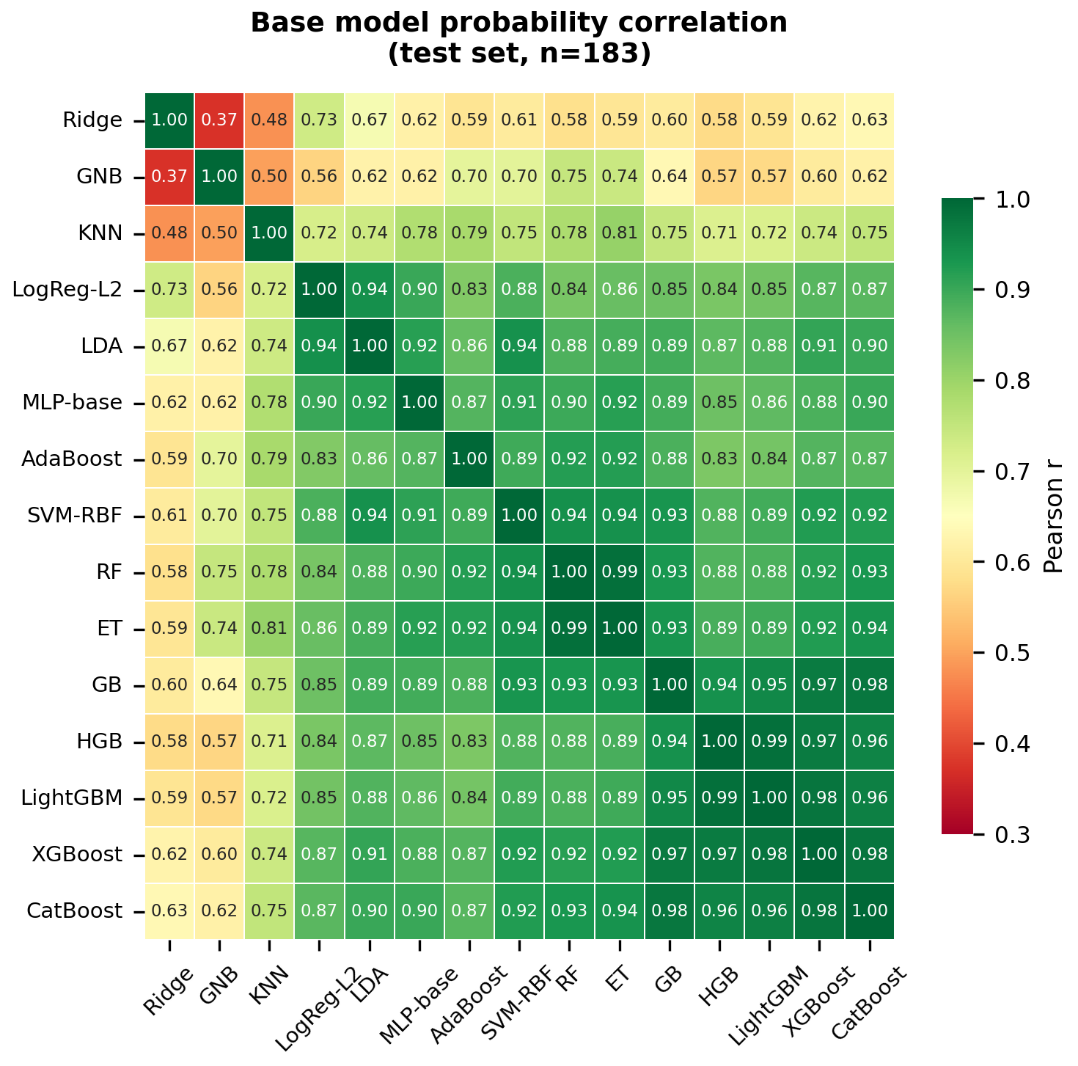

Supplementary Figure S4. Correlation among base-model probability outputs. Pearson correlation matrix of independent test-set probability outputs from the 15 base classifiers. The matrix shows shared and partially distinct prediction patterns among base learners, supporting the use of stacked integration.

Supplementary Table S1. Definitions and equations of custom RNA sequence complexity, repeat, motif and predicted-structure descriptors.

| Feature name / pattern | Feature group | Definition | Readable equation / calculation |
| --- | --- | --- | --- |
| length | Sequence length | Total number of canonical RNA nucleotides after preprocessing. | L = number of nucleotides in the cleaned RNA sequence |
| entropy_1mer | Sequence entropy | Sequence entropy calculated from single-nucleotide frequencies. | E1mer = − Σ pi log2(pi), where pi is the frequency of each nucleotide: A, U, G or C. |
| entropy_2mer | Sequence entropy | Sequence entropy calculated from dinucleotide frequencies. | E2mer = − Σ pi log2(pi), where pi is the frequency of each dinucleotide. |
| entropy_3mer | Sequence entropy | Sequence entropy calculated from trinucleotide frequencies. | E3mer = − Σ pi log2(pi), where pi is the frequency of each trinucleotide. |
| linguistic_complexity_k1_5 | Sequence complexity | Mean observed-to-possible k-mer ratio across k-mer lengths from 1 to 5. | LC = 1/5 Σk=1 to 5 [ Uk / min(4k, L − k + 1) ], where Uk is the number of unique k-mers observed at length k, and L is the RNA sequence length. |
| low_complexity_burden_entropy12 | RNA nucleotide-level low-complexity burden | Fraction of 12-nt windows with sequence entropy below 1.2. This refers to RNA sequence-level low complexity, not protein low-complexity domains. | LCBentropy12 = N(Ew < 1.2) / Nw, where Ew is the sequence entropy of window w, and Nw is the total number of 12-nt windows. |
| low_complexity_burden_unique12 | RNA nucleotide-level low-complexity burden | Fraction of 12-nt windows containing two or fewer unique nucleotide types. | LCBunique12 = N(Uw ≤ 2) / Nw, where Uw is the number of unique nucleotide types in window w, and Nw is the total number of 12-nt windows |
| compression_ratio_gzip | Sequence compressibility | Ratio of gzip-compressed sequence length to original sequence length. | CRgzip = Lcompressed / Lraw, where Lcompressed is the gzip-compressed sequence length and Lraw is the original RNA sequence length. |
| A_run_max, U_run_max, G_run_max, C_run_max | Homopolymer runs | Longest uninterrupted run of a specific nucleotide. | RunMaxb = maximum length of a continuous run of nucleotide b, where b is A, U, G or C. |
| run_max_any | Homopolymer runs | Longest uninterrupted homopolymer run across all nucleotide types. | RunMaxany = maximum of A_run_max, U_run_max, G_run_max and C_run_max |
| A_run_count_ge4, U_run_count_ge4, G_run_count_ge4, C_run_count_ge4 | Homopolymer runs | Number of nucleotide-specific homopolymer runs with length ≥4. | RunCountb≥4 = number of continuous runs of nucleotide b with length ≥4 |
| A_run_count_ge4_per100nt, U_run_count_ge4_per100nt, G_run_count_ge4_per100nt, C_run_count_ge4_per100nt | Homopolymer runs | Number of nucleotide-specific homopolymer runs of length ≥4, normalized per 100 nt. | RunCountb≥4 per 100 nt = (RunCountb≥4 / L) × 100 |
| homopolymer_coverage_ge4 | Homopolymer coverage | Fraction of the RNA sequence covered by homopolymer runs of length ≥4. | Homopolymer coverage≥4 = total length of all homopolymer runs ≥4 / L |
| tandem_repeat_density_m3 | Tandem-repeat organization | Fraction of the sequence covered by trinucleotide tandem-repeat regions. At least three consecutive repeat units were required. | TR density m3 = length covered by 3-nt tandem repeats / L |
| tandem_repeat_regions_m1, tandem_repeat_regions_m2, tandem_repeat_regions_m3 | Tandem-repeat organization | Number of separate tandem-repeat regions detected for motif lengths of 1, 2 or 3 nt. | TR regions m = number of separate tandem-repeat regions with motif length m, where m = 1, 2 or 3. |
| tandem_repeat_longest_region_m1, tandem_repeat_longest_region_m2, tandem_repeat_longest_region_m3 | Tandem-repeat organization | Length of the longest tandem-repeat region for motif lengths of 1, 2 or 3 nt. | TR longest region m = maximum length of a tandem-repeat region with motif length m |
| tandem_repeat_max_repeat_count_m1, tandem_repeat_max_repeat_count_m2, tandem_repeat_max_repeat_count_m3 | Tandem-repeat organization | Maximum number of consecutive repeat units observed in a tandem-repeat region. | TR max repeat count m = maximum number of consecutive repeat units for motif length m |
| UG_density, GG_density, CC_density, UA_density, AU_density, UU_density, CUG_density, AUA_density, UAU_density, UAC_density | Motif density | Frequency of a predefined RNA motif normalized by the number of possible motif positions. | Motif densityx = Nx / (L − motif length + 1), where Nx is the number of occurrences of motif x. |
| UG_cluster_max30, GG_cluster_max30, CC_cluster_max30, UA_cluster_max30, AU_cluster_max30, UU_cluster_max30, CUG_cluster_max30, AUA_cluster_max30, UAU_cluster_max30, UAC_cluster_max30 | Motif clustering | Maximum number of occurrences of a predefined motif within any 30-nt window. | Motif clusterx,30 = maximum number of motif x occurrences in any 30-nt window |
| mfe | Predicted secondary structure | Minimum free energy of the RNA secondary structure predicted using RNAfold/ViennaRNA. | MFE = minimum free energy predicted by RNAfold |
| mfe_per_nt | Predicted secondary structure | Minimum free energy normalized by RNA sequence length. | MFE per nt = MFE / L |
| paired_fraction | Predicted secondary structure | Fraction of nucleotides predicted to be base-paired in the RNAfold dot-bracket structure. | Paired fraction = number of paired nucleotides / L |
| max_stem_len | Predicted secondary structure | Maximum length of a continuous paired stem segment in the predicted secondary structure. | Max stem length = longest continuous paired stem segment |
| max_unpaired_run | Predicted secondary structure | Longest continuous unpaired region in the predicted secondary structure. | Max unpaired run = longest continuous unpaired segment |

Supplementary Table S2. Handcrafted RNA feature families and calculation tools. Standard composition, k-mer and DACC descriptors are calculated using iLearn. Additional repeat, motif density, structure and complexity descriptors are calculated using custom Python scripts or RNAfold-derived outputs.

| **Feature family** | **Features** | **Description** | **Computed using / source** | **Number retained** |
| --- | --- | --- | --- | --- |
| Mononucleotide frequency | A, G | Nucleotide composition features; U and C omitted to avoid perfect collinearity | iLearn | 2 |
| Dinucleotide frequency | dinucleotide features(AG, AU, CA, CC, CU..) | Selected dinucleotide composition features after redundancy reduction | iLearn | 13 |
| Composition / skew | GC_skew, AU_skew, purine_content, amino_content | Global nucleotide composition and compositional imbalance | Custom Python sequence-statistics script | 4 |
| Trinucleotide frequency | trinucleotide features (CUA, UAA, AUA, AAU, UAU..) | Selected 3-mer/k-mer frequency features after redundancy reduction | iLearn | 47 |
| DACC descriptors | DACC features (dacc_1, dacc_2, dacc_3, …) | Dinucleotide auto-cross-covariance descriptors capturing long-range sequence-order information | iLearn | 63 |
| Lambda autocorrelation | lambda_1, lambda_2 | Autocorrelation descriptors summarizing sequence-order dependence | iLearn / custom post-processing if applicable | 2 |
| Homopolymer / run-length | U/A/G/C run features, run_max_any, homopolymer_coverage_ge4 | Homopolymer counts, maximum run lengths, and run coverage | Custom Python regular-expression/run-length parser | 11 |
| Motif cluster density | AU_cluster_max30, CUG_cluster_max30, GG_cluster_max30, UAC_cluster_max30, UAU_cluster_max30, UU_cluster_max30 | Maximum motif count within any 30-nt sliding window | Custom Python sliding-window motif counter | 6 |
| Motif linear density | CUG_density, UAU_density | Global motif density per nucleotide | Custom Python motif counter | 2 |
| Tandem repeat statistics | tandem repeat density/count/region features | Tandem-repeat density and repeat-region descriptors for motif lengths 1–3 | Custom Python tandem-repeat scanning script | 6 |
| Predicted RNA secondary structure | mfe, mfe_per_nt, paired_fraction, max_stem_len, max_unpaired_run | Minimum-free-energy folding and derived structural descriptors | ViennaRNA RNAfold + custom Python parser | 5 |
| Sequence complexity / entropy | linguistic_complexity_k1_5, entropy_1mer, entropy_2mer, compression_ratio_gzip, low_complexity_burden_unique12 | Sequence complexity, Shannon entropy, compressibility, and repetitive k-mer burden | Custom Python entropy/complexity scripts + Python gzip | 5 |
| Sequence length | length | Total sequence length in nucleotides | Custom Python sequence parser | 1 |

Supplementary Table S4. Hyperparameters of the base learners and stacked Meta-NN. Scale-normalized models trained inside a Standard Scalar pipeline; tree-based models trained on unscaled features.

| **Slno** | **Model** | **Family** | **Hyperparameters** | **Scaled** |
| --- | --- | --- | --- | --- |
| 1 | Logistic Regression | Linear classifier | Penalty = L2; C = 1.0; solver = lbfgs; max_iter = 4000; random_state = 42 | ✓ |
| 2 | Ridge Classifier | Regularized linear classifier | alpha = 1.0; solver = lsqr; random_state = 42 | ✓ |
| 3 | LDA | Linear discriminant analysis | solver = lsqr; shrinkage = auto | ✓ |
| 4 | KNN | Distance-based classifier | n_neighbors = 15; metric = Minkowski/L2; weights = uniform; n_jobs = −1 | ✓ |
| 5 | Gaussian NB | Probabilistic classifier | var_smoothing = 1e−7 | ✓ |
| 6 | Random Forest | Tree ensemble | n_estimators = 500; criterion = Gini; max_features = sqrt; max_depth = None; min_samples_split = 2; min_samples_leaf = 1; bootstrap = True; n_jobs = −1; random_state = 42 | — |
| 7 | Extra Trees | Tree ensemble | n_estimators = 500; criterion = Gini; max_features = sqrt; max_depth = None; splitter = random; bootstrap = False; n_jobs = −1; random_state = 42 | — |
| 8 | AdaBoost | Boosting ensemble | n_estimators = 300; learning_rate = 1.0; base estimator = decision stump; algorithm = SAMME.R; random_state = 42 | — |
| 9 | Gradient Boosting | Boosting ensemble | n_estimators = 300; learning_rate = 0.05; max_depth = 3; min_samples_split = 2; min_samples_leaf = 1; subsample = 1.0; loss = log loss; random_state = 42 | — |
| 10 | Hist Gradient Boosting | Boosting ensemble | max_iter = 500; learning_rate = 0.05; max_depth = None; max_leaf_nodes = 31; min_samples_leaf = 20; l2_regularization = 0.0; max_bins = 255; early stopping = auto; random_state = 42 | — |
| 11 | LightGBM | Gradient-boosted trees | n_estimators = 500; learning_rate = 0.05; num_leaves = 63; min_child_samples = 20; subsample = 0.9; colsample_bytree = 0.9; boosting type = GBDT; objective = binary; n_jobs = −1; random_state = 42 | — |
| 12 | XGBoost | Gradient-boosted trees | n_estimators = 500; learning_rate = 0.05; max_depth = 6; subsample = 0.9; colsample_bytree = 0.9; tree_method = hist; eval_metric = logloss; objective = binary:logistic; n_jobs = −1; random_state = 42 | — |
| 13 | CatBoost | Gradient-boosted trees | iterations = 500; learning_rate = 0.05; depth = 6; l2_leaf_reg = 3.0; loss function = Logloss; bootstrap type = Bayesian; verbose = False; allow_writing_files = False; random_seed = 42 | — |
| 14 | SVM-RBF | Kernel classifier | C = 1.0; kernel = RBF; gamma = scale; probability = True; random_state = 42 | ✓ |
| 15 | MLP-base | Neural network | hidden layers = (128, 64); activation = ReLU; solver = Adam; alpha = 1e−3; batch size = auto; learning_rate_init = 0.001; max_iter = 300; early stopping = True; validation_fraction = 0.15; n_iter_no_change = 10; random_state = 42 | ✓ |
| 16 | **Stacked Meta-NN** | **Neural meta-classifier** | Input = 129 dimensions: 50 tabular PCs + 64 RNA-FM PCs + 15 base-model probabilities; hidden layers = (64, 16); activation = ReLU; solver = Adam; alpha = 1e−3; batch_size = 64; learning_rate_init = 1e−3; max_iter = 500; early stopping = True; validation_fraction = 0.20; n_iter_no_change = 25; output = sigmoid probability; 25-seed probability averaging | ✓ |

Supplementary Table S5. Model evaluation metrics and definitions.

| **Metric** | **Equation** | **Interpretation** |
| --- | --- | --- |
| Accuracy | (TP + TN) / (TP + TN + FP + FN) | Overall fraction of correctly classified RNAs |
| Precision | TP / (TP + FP) | Fraction of predicted LLPS+ RNAs that are true LLPS+ |
| Recall / Sensitivity | TP / (TP + FN) | Fraction of true LLPS+ RNAs correctly identified |
| Specificity | TN / (TN + FP) | Fraction of LLPS− RNAs correctly identified |
| F1 score | 2 × Precision × Recall / (Precision + Recall) | Harmonic mean of precision and recall |
| MCC | (TP × TN − FP × FN) / √[(TP + FP)(TP + FN)(TN + FP)(TN + FN)] | Balanced classification metric using all confusion-matrix terms |
| False positive rate | FP / (FP + TN) = 1 − Specificity | Fraction of LLPS− RNAs incorrectly predicted as LLPS+ |
| AUC-ROC | Area under the receiver operating characteristic curve | Threshold-independent ranking performance based on TPR and FPR |
| AUC-PR | Area under the precision–recall curve | Threshold-independent ranking performance based on precision and recall |

Supplementary Table S6. Representative sequence-derived features shown in Figures 2 and 3. Selected handcrafted descriptors plotted in the main feature-distribution figures. Features were selected from the training-set univariate analysis to represent major sequence-feature families, including composition/base skew, local k-mer grammar, sequence entropy, low-complexity burden, long-range dinucleotide autocorrelation, repeat organization and predicted secondary structure. q-values are calculated using two-sided Mann–Whitney U tests followed by Benjamini–Hochberg FDR correction. Effect sizes are reported as rank-biserial effect sizes. Positive values indicate higher feature values in LLPS+ RNAs, whereas negative values indicate higher feature values in LLPS− background RNAs.

| **Feature group** | **Feature** | **Direction in LLPS+** | **q-value from S3** | **Effect size from S3** |
| --- | --- | --- | --- | --- |
| Composition | amino_content | Lower | 2.00e−11 | −0.302 |
| Composition | AU_skew | Lower | 1.95e−05 | −0.194 |
| Homopolymer run | A_run_count_ge4_per100nt | Lower | 9.36e−14 | −0.331 |
| k-mer grammar | CUA | Lower | 3.87e−28 | −0.489 |
| k-mer grammar | UAA | Lower | 1.25e−25 | −0.465 |
| k-mer grammar | ACU | Lower | 2.23e−25 | −0.461 |
| k-mer grammar | UA | Lower | 1.81e−18 | −0.393 |
| k-mer grammar | AUA | Lower | 3.31e−18 | −0.389 |
| k-mer grammar | AAU | Lower | 6.63e−15 | −0.348 |
| Low-complexity | low_complexity_burden_unique12 | Higher | 2.53e−07 | +0.232 |
| Long-range composition | dacc_57 | Higher | 8.95e−07 | +0.222 |
| Long-range composition | dacc_20 | Higher | 7.30e−06 | +0.204 |
| Long-range composition | dacc_39 | Higher | 2.77e−05 | +0.190 |
| Long-range composition | dacc_8 | Higher | 1.63e−04 | +0.172 |
| Tandem / repeat | tandem_repeat_max_repeat_count_m3 | Higher* | 7.65e−06 | +0.193 |
| Tandem / repeat | tandem_repeat_regions_m3 | Higher* | 1.96e−04 | +0.163 |
| Predicted structure | max_stem_len | Lower | 7.64e−02 | −0.084 |
| Predicted structure | mfe_per_nt | Limited separation | 8.59e−01 | −0.010 |
| Predicted structure | paired_fraction | Limited separation | 6.83e−01 | +0.021 |
| Predicted structure | max_unpaired_run | Higher | 2.29e−01 | +0.058 |

Note: Direction in LLPS+ is assigned according to the sign of the rank-biserial effect size rather than median difference alone. Thus, some features, particularly sparse repeat-related descriptors, may have equal class medians but still show a positive or negative rank-biserial effect size because of differences in the full distribution. In such cases, “Higher” indicates an overall positive distributional shift in LLPS+ RNAs.

Supplementary Table S7. Predictive performance of logistic-regression baselines across feature representations. All models use the same L2-regularized logistic-regression pipeline and differ only in their input features. Bold values indicate the best performance for each metric.

| Feature representation | No. of features | Accuracy | F1 | AUC-ROC | AUC-PR | MCC |
| --- | --- | --- | --- | --- | --- | --- |
| Length + GC only | 2 | 0.574 | 0.316 | 0.469 | 0.562 | 0.147 |
| Handcrafted only | 167 | 0.820 | 0.800 | 0.879 | 0.895 | 0.638 |
| RNA-FM only | 640 | 0.847 | 0.833 | 0.909 | 0.908 | 0.693 |
| Handcrafted + RNA-FM | 807 | 0.869 | 0.860 | 0.924 | 0.929 | 0.737 |

Supplementary Table S8. Bootstrap confidence intervals for the final stacked predictor. Confidence intervals were estimated from 2,000 bootstrap resamples of the independent test-set predictions.

| **Metric** | **Point estimate** | **95% CI lower** | **95% CI upper** |
| --- | --- | --- | --- |
| Accuracy | 0.8907 | 0.8415 | 0.9344 |
| Precision | 0.9125 | 0.8452 | 0.9706 |
| Recall | 0.8488 | 0.7701 | 0.9221 |
| F1 | 0.8795 | 0.8212 | 0.9282 |
| AUC-ROC | 0.9348 | 0.8928 | 0.9676 |
| AUC-PR | 0.9443 | 0.9066 | 0.9712 |
| MCC | 0.7815 | 0.6847 | 0.8682 |

Supplementary Table S9. Ablation analysis of stacked predictor design choices. Each configuration is evaluated on the independent test set. Bold values indicate the best value for each metric.

| **Config** | **Description** | **Meta-model input** | **Dimension** | **Accuracy** | **F1** | **AUC-ROC** | **AUC-PR** | **MCC** |
| --- | --- | --- | --- | --- | --- | --- | --- | --- |
| A | XGBoost only, no stacking | — | — | 0.863 | 0.850 | 0.919 | 0.932 | 0.726 |
| B | Stacking with base probabilities only | Base probabilities | 15 | 0.885 | 0.874 | 0.932 | 0.940 | 0.770 |
| C | Stacking + tabular PCA | Tabular PCs + base probabilities | 65 | 0.880 | 0.869 | 0.928 | 0.938 | 0.759 |
| D | Stacking + RNA-FM PCA | RNA-FM PCs + base probabilities | 79 | 0.880 | 0.868 | **0.937** | 0.943 | 0.759 |
| E | Stacking + joint PCA(64) | Joint PCs + base probabilities | 79 | 0.885 | 0.874 | 0.933 | 0.941 | 0.770 |
| F | Stacking + separate PCA | Tabular PCs + RNA-FM PCs + base probabilities | 129 | **0.891** | **0.880** | 0.935 | **0.944** | **0.782** |
| G | Stacking + joint PCA(114) | Joint PCs + base probabilities | 129 | 0.842 | 0.824 | 0.912 | 0.902 | 0.680 |
| H | Base models trained on PCA features | Base probabilities only | 15 | 0.847 | 0.827 | 0.902 | 0.885 | 0.692 |

Supplementary Table S10. Top SHAP-ranked features contributing to LLPS-associated RNA prediction. SHAP values are calculated using a LightGBM model trained on the full 807-feature representation. Features are ranked by mean absolute SHAP value, which reflects global contribution to model predictions. “Low → LLPS+” indicates that lower feature values increased the predicted probability of the LLPS+ class, whereas “High → LLPS+” indicates that higher feature values increased the predicted probability of the LLPS+ class. SHAP direction reflects model behaviour and may differ from univariate class-wise feature differences.

| **Rank** | **Feature** | **Feature type** | **Mean \|SHAP\|** | **Feature-value effect on prediction** |
| --- | --- | --- | --- | --- |
| 1 | linguistic_complexity_k1_5 | Handcrafted | 1.846 | Low → LLPS+ |
| 2 | CUU | Handcrafted | 0.682 | Low → LLPS+ |
| 3 | RNA-FM dim 413 | RNA-FM | 0.512 | High → LLPS+ |
| 4 | CUA | Handcrafted | 0.505 | Low → LLPS+ |
| 5 | RNA-FM dim 563 | RNA-FM | 0.469 | Low → LLPS+ |
| 6 | RNA-FM dim 100 | RNA-FM | 0.434 | Low → LLPS+ |
| 7 | RNA-FM dim 284 | RNA-FM | 0.404 | High → LLPS+ |
| 8 | AUU | Handcrafted | 0.363 | Low → LLPS+ |
| 9 | RNA-FM dim 426 | RNA-FM | 0.341 | High → LLPS+ |
| 10 | UAU_density | Handcrafted | 0.332 | High → LLPS+ |
| 11 | dacc_21 | Handcrafted | 0.301 | High → LLPS+ |
| 12 | RNA-FM dim 34 | RNA-FM | 0.283 | Low → LLPS+ |
| 13 | UAA | Handcrafted | 0.282 | High → LLPS+ |
| 14 | RNA-FM dim 280 | RNA-FM | 0.255 | High → LLPS+ |
| 15 | dacc_24 | Handcrafted | 0.241 | Low → LLPS+ |
| 16 | RNA-FM dim 134 | RNA-FM | 0.236 | Low → LLPS+ |
| 17 | U_run_count_ge4 | Handcrafted | 0.200 | Low → LLPS+ |
| 18 | RNA-FM dim 577 | RNA-FM | 0.199 | Low → LLPS+ |
| 19 | RNA-FM dim 528 | RNA-FM | 0.189 | High → LLPS+ |
| 20 | RNA-FM dim 380 | RNA-FM | 0.186 | Low → LLPS+ |

Supplementary Table S11. Feature-family-level summary of handcrafted RNA descriptors. Feature families are summarized using univariate comparisons between LLPS− background RNAs and LLPS+ reviewed RNAs. Significant features are defined using Benjamini–Hochberg FDR correction. Effect size refers to absolute rank-biserial correlation.

| **Feature family** | **Total features** | **Significant features** | **Significant (%)** | **Features passing q + effect threshold** | **Median \|effect size\|** | **Maximum \|effect size\|** | **Features higher in LLPS+** | **Features higher in LLPS−** | **Dominant direction** |
| --- | --- | --- | --- | --- | --- | --- | --- | --- | --- |
| Composition / k-mer | 66 | 55 | 83.3 | 47 | 0.236 | 0.489 | 3 | 63 | Higher in LLPS− |
| DACC descriptors | 63 | 29 | 46.0 | 13 | 0.084 | 0.229 | 37 | 26 | Higher in LLPS+ |
| Motif clustering | 8 | 6 | 75.0 | 3 | 0.129 | 0.270 | 1 | 7 | Higher in LLPS− |
| Homopolymer / run | 11 | 4 | 36.4 | 1 | 0.066 | 0.331 | 5 | 6 | Mixed / higher in LLPS− |
| Tandem / repeat | 6 | 5 | 83.3 | 2 | 0.117 | 0.193 | 6 | 0 | Higher in LLPS+ |
| Low-complexity | 5 | 3 | 60.0 | 3 | 0.187 | 0.253 | 2 | 3 | Mixed |
| Other derived | 2 | 1 | 50.0 | 0 | 0.057 | 0.112 | 2 | 0 | Higher in LLPS+ |
| Predicted secondary structure | 5 | 0 | 0.0 | 0 | 0.033 | 0.084 | 2 | 3 | Weak / not significant |
| Length | 1 | 0 | 0.0 | 0 | 0.041 | 0.041 | 0 | 1 | Not significant |
| Total | 167 | 103 | — | 69 | — | — | — | — | — |

**Note:**

Positive effect sizes indicate higher feature values in LLPS+ RNAs, whereas negative effect sizes indicate lower feature values in LLPS+ RNAs. “Dominant direction” summarizes the overall direction of significant or non-negligible effects within each feature family.
